# Trim-Away degrades lysine-less substrates without requiring ligase autoubiquitination

**DOI:** 10.1101/2022.06.29.498105

**Authors:** Leo Kiss, Tyler Rhinesmith, Claire F Dickson, Jonas Weidenhausen, Shannon Smyly, Jakub Luptak, Ji-Chun Yang, Sarah L Maslen, Irmgard Sinning, David Neuhaus, Dean Clift, Leo C James

**Affiliations:** MRC Laboratory of Molecular Biology, Francis Crick Avenue, Cambridge CB2 0QH, United Kingdom; EMBL Australia Node in Single Molecule Science and ARC Centre of Excellence in Advanced Molecular Imaging School of Medical Sciences, UNSW Sydney, NSW, 2052, Australia; Biochemiezentrum der Universität Heidelberg (BZH), INF328, D-69120 Heidelberg, Germany; Max Planck Institute of Biochemistry, AM Klopferspitz 18, 82152 Martinsried, Germany; EMBL Heidelberg, Meyerhofstraße 1, 69117 Heidelberg, Germany; The Francis Crick Institute, 1 Midland Road, London NW1 1AT, United Kingdom

**Author notes:** Correspondence: Leo Kiss; Dean Clift; Leo C. James.

## Abstract

TRIM proteins are the largest family of E3 ligases in mammals. They include the intracellular antibody receptor TRIM21, which is responsible for mediating targeted protein degradation during Trim-Away. Despite their importance, the ubiquitination mechanism of TRIM ligases has remained elusive. Here we show that while TRIM21 activation results in ubiquitination of both ligase and substrate, autoubiquitination is regulatory and not required for substrate degradation. Substrate binding stimulates N-terminal RING autoubiquitination by the E2 Ube2W, but when inhibited by N-terminal acetylation this prevents neither substrate ubiquitination nor degradation and has no impact on TRIM21 antiviral activity. Instead, uncoupling ligase and substrate degradation prevents ligase recycling and extends functional persistence in cells. Substrate ubiquitination mediates degradation but Trim-Away efficiently degrades lysine-less substrates, suggesting a non-canonical ubiquitination mechanism explains its broad substrate specificity.

## INTRODUCTION

TRIM proteins constitute the largest family of RING E3 ligases in mammals. They include TRIMs that suppress viral infection (TRIM5^1^, TRIM21^2^, TRIM22^3^, TRIM25^4^), activate innate immunity (TRIM32^5^, TRIM56^6^, TRIM65^7^, RIPLET^8^), and repress transcription (TRIM4^9^, TRIM28^10^). Unlike cullin-RING ubiquitin ligases (CRLs), which use a modular system of RINGs, adaptors and scaffolds assembled combinatorially to create distinct enzymes^11,12^, TRIM ligases contain both substrate-targeting and catalytic domains in one polyprotein. How TRIMs catalyse ubiquitination is incompletely understood, particularly in terms of activation, ubiquitin priming and chain extension. This is due in part to the difficulty linking in vitro activity with cellular function. For instance, many RINGs have been shown to work with E2s in vitro for which there is no data supporting a cellular role.

Current mechanisms of TRIM catalysis have been informed primarily by experiments on the two antiviral proteins TRIM5 and TRIM21. Both proteins are dimers containing a RING, B Box, coiled-coil and PRYSPRY domains. Each RING domain is arranged at opposite ends of the elongated antiparallel coiled-coil^13^ and whilst ubiquitination of monomeric RINGs can be detected in vitro, dimerization is required for full cellular activity^14,15^. Intramolecular RING dimerization would require an extensive conformational rearrangement, involving an extreme bend angle in the coiled-coil, and existing data suggests that TRIM RINGs instead undergo intermolecular dimerization through a mechanism of substrate-induced clustering^15^. In the case of TRIM5, this occurs during binding to the conical capsid of HIV-1^16^: The primarily hexameric capsid induces formation of a hexameric lattice of TRIM5 molecules^17^ anchored to the capsid surface through PRYSPRY domain interactions^16^. The TRIM5 lattice is further stabilized through trimeric contacts formed between the B Box domains at each vertex^18^ and transient RING dimerization^18,19^. TRIM21 also undergoes supramolecular clustering^15^, including on the surface of viral capsids^20^, but is anchored to its substrates by an intermediate antibody molecule^2^: The Fabs of each antibody bind the substrate whilst the Fc is bound by the TRIM21 PRYSPRY^21^. There is no evidence that TRIM21 forms a regular structure or that its B Box mediates oligomerization. Instead, the B Box of TRIM21 is an autoinhibitory domain that supresses RING activity in the non-clustered state by competing for E2∼Ub binding^14^. Supramolecular assembly is sufficient for TRIM RING activation. In the case of TRIM21, light-induced clustering of a cryptochrome2-TRIM21 fusion triggered its TRIM21 and proteasome-dependent degradation^15^. Meanwhile, TRIM5 degradation was accelerated in the presence of HIV-1 capsid^22^, and prevented by a single B Box mutation that prevents higher order assembly^23^.

A further important difference between TRIM and CRL ligases is that the former undergoes degradation along with its substrate. This has been shown for TRIM5 during HIV infection^22^ and for TRIM21 with a wide-range of substrates during Trim-Away^24^. Moreover, TRIM21 and its substrates are degraded with matching kinetics suggesting that they are processed together as a complex^24^. In support of TRIM ligase self-degradation, light-induced clustering of a TRIM21 RING-crytochrome2 fusion was sufficient to cause ligase degradation^15^. Meanwhile, TRIM5 self-degradation can be induced simply by ectopic overexpression^25^, which leads to the formation of large oligomers called ‘cytoplasmic bodies’, likely driven by B Box trimerization^18,23^.

Activation of TRIM RINGs enables their recruitment of E2s and catalysis of ubiquitination. Multiple E2s have been reported as partners but only depletion of the N-terminal monoubiquitinating E2 Ube2W or the K63-chain forming heterodimer Ube2N/2V2 has been shown to inhibit the cellular function of TRIM5^26,27^ and TRIM21^28,29^. Moreover, point mutations in TRIM21, which specifically inhibit catalysis with Ube2N resulted in loss of cellular function^15,30^. K63-chain ubiquitination has also been implicated in the function of many other TRIMs, such as TRIM4^31^, TRIM8^32^, TRIM22^33^, TRIM31^34^, TRIM34^35^, TRIM54^36^, TRIM59^37^, TRIM65^7^ and RIPLET^8^. In vitro, both TRIM5 and TRIM21 have been shown to catalyse monoubiquitination of their own N-terminus when incubated with Ube2W, whilst the addition of Ube2N/2V2 drives chain extension to produce an anchored K63 chain^26,28^. This K63-linked autoubiquitination can be detected in cells during substrate engagement^15^ or over-expression and is reversed by Ube2W or Ube2N depletion^26,28^. Substrate-induced RING activation can also be reproduced in vitro, with the addition of IgG Fc promoting the formation of anchored K63-chains on TRIM21^38^. Moreover, in vitro experiments suggest that RING clustering is important not only to generate active RING dimers but also to allow intermolecular RING ubiquitination. A ‘two-plus-one’ model has been demonstrated, in which a RING dimer ubiquitinates the N-terminus of a neighbouring RING monomer^19^. For TRIM5, this is supported by in vitro ubiquitination rescue experiments using catalytically dead mutants^19^ and structural data demonstrating a trimeric RING arrangement in assembled TRIM5 lattices^18,39,40^. For TRIM21, RING dimerization was shown to be insufficient for effective substrate-induced ubiquitination, with full activity requiring the recruitment of two RING dimers^38^.

This correspondence between the functional requirement for TRIM clustering and the mechanistic requirement for higher-order catalytic RING topology helps explain how TRIMs are activated and regulated. However, how K63-ubiquitination facilitates proteasomal degradation is less clear. It has been suggested that TRIM-synthesized K63-chains are further modified with branched K48-chains, similar to that reported for UBR5/HUWE1/UBR4^41,42^. In support of this, both K63 and K48-chains can be detected on overexpressed TRIM21 and are lost concomitantly upon either Ube2W or Ube2N depletion^28^. Importantly, while autoubiquitination explains why TRIM ligases are degraded upon activation, it does not provide a direct mechanism for substrate degradation. Whether substrates are also modified with ubiquitin during their cellular degradation is unknown, despite attempts to detect this^2,16^, and so because of the absence of such data it has been proposed that ligase autoubiquitination alone may drive proteasome recruitment, resulting in degradation of the entire TRIM:substrate complex^2,25,43,44^.

Here we sought to test the requirement for TRIM autoubiquitination in substrate degradation. Using TRIM21 as a model system, we show that whilst N-terminal ubiquitination drives ligase turnover it is not required for substrate degradation. Rather, uncoupling ligase and substrate degradation prolongs ligase lifetime allowing it to persist in cells for longer. We demonstrate, both in vitro and in cells, direct substrate ubiquitination by TRIM21 and show that this is unaffected by inhibiting N-terminal TRIM21 autoubiquitination. Finally, we establish a cellular degradation assay in which all lysines can be removed or mutated to arginine. Surprisingly, we find that degradation does not require lysines in either ligase or substrate, suggesting that either N-terminal substrate modification or non-canonical ubiquitination is required for TRIM21 mediated degradation.

## RESULTS

### Structure of TRIM21 RING in complex with Ube2W

To understand how TRIM21 recruits Ube2W, we solved the crystal structure of the RING domain in complex with a Ube2W dimerization mutant^45^ in which the active site cysteine was also replaced with lysine (Ube2W^V30K/D67K/C91K^). Two copies of each Ube2W and RING could be found in the asymmetric unit, with the two RINGs forming a homodimer as described previously^14,30,38^ (Figure 1A). RING and Ube2W engage each other via the canonical RING:E2 interface (Figure 1B). By superposing the RING:Ube2W structure with Ube2N∼Ub from the previously determined Ub-RING:Ube2N∼Ub:Ube2V2 structure^38^, the activation of Ube2W∼Ub was modelled (Figure 1C). Overall, the arrangement of a Ube2W∼Ub is similar to Ube2N∼Ub, when being activated by TRIM21^30,38^. In this model, the donor ubiquitin is in the closed conformation, stabilized by both RING protomers. Interestingly, the model also suggests that TRIM21 E13 might engage ubiquitin K11 to stabilize the closed conformation (Figure 1D). E13 is part of a tri-ionic motif that was identified to drive Ube2N∼Ub interaction ^30^. We tested whether this motif is involved in Ube2W binding by performing NMR titrations of ^15^N-labelled TRIM21 tri-ionic mutants against Ube2W^V30K/D67K/C91K^. Mutation of the tri-ionic residues E12 and E13 to alanine did not lead to obvious reductions in the observed chemical shift perturbations (CSPs) (Supplementary Figure 1A-C). Moreover, tri-ionic mutants had only a modest effect on TRIM21 monoubiquitination (Supplementary Figure 1D). TRIM21 mutants E13A and E13R both showed a slight reduction in activity, suggesting that residue E13 could indeed interact with ubiquitin K11, but this is not as critical as for Ube2N∼Ub^30^. When comparing the RING in the RING:Ube2W structure to the apo-^14^ and Ube2N∼Ub^30^ engaged structures, we noted that the N- and C-terminal helices of the RINGs are partly unfolded when bound by Ube2W (Supplementary Figure 1E). While this may reflect differences in crystallization, it could suggest that Ube2W has the potential to destabilize the 4-helix bundle, thereby generating a disordered N-terminus for modification. Even so, this is insufficient to explain how Ube2W can modify the N-terminus of a RING it is bound to, as it is located far away from the E2 active site.

**Figure 1:**
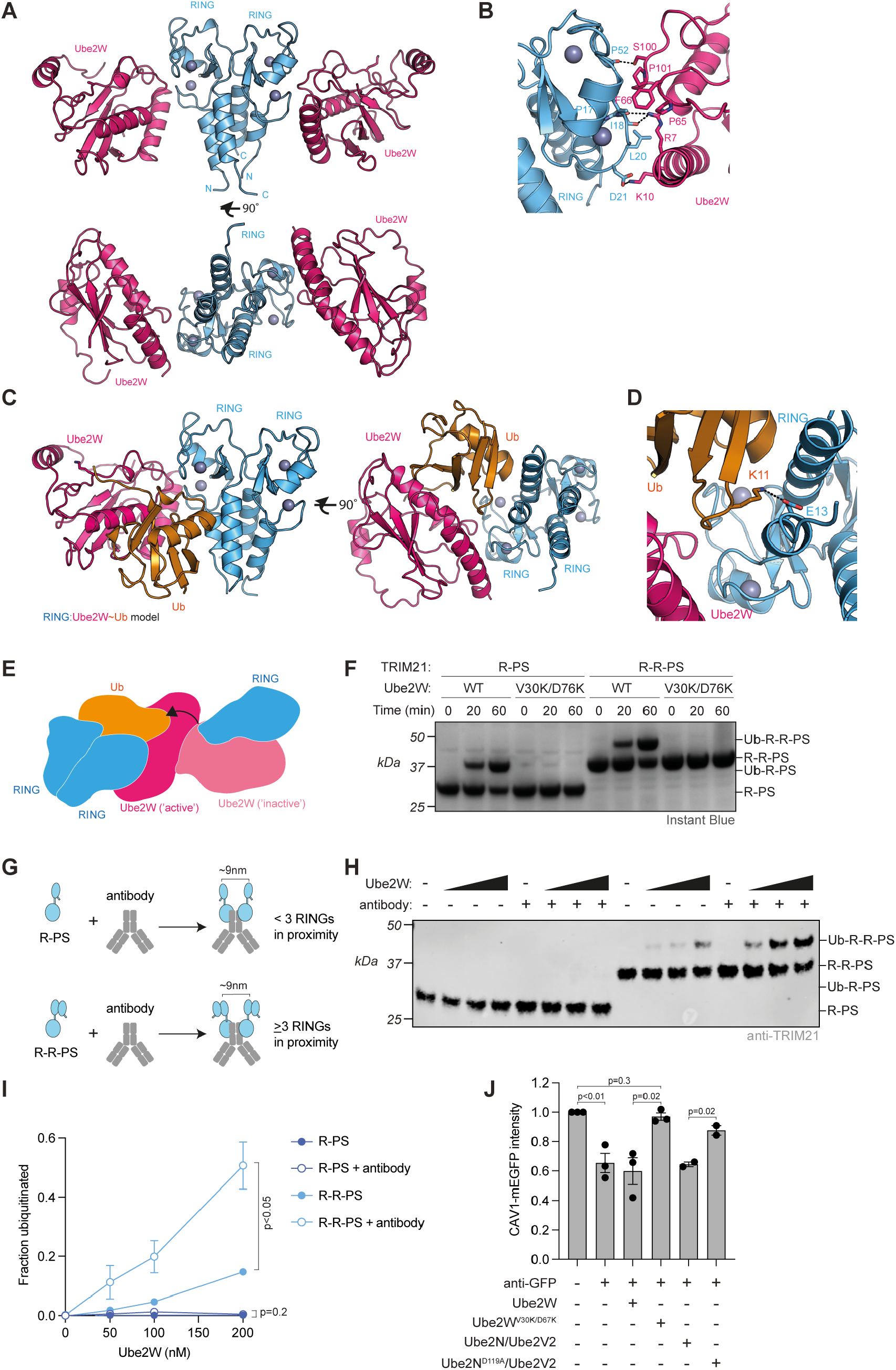
Dimeric Ube2W and RING clustering is required for ligase autoubiquitination. **(A)** 2.25 Å X-ray structure of TRIM21 RING (blue) in complex with Ube2W (pink). (**B)** Close-up of the E2:E3 interface. (**C**) Structural model of a RING:Ube2W∼Ub complex based on superposition of the RING:Ube2W structure and the Ub-RING:Ube2N∼Ub:Ube2V2 structure (7BBD)^38^. Ube2N∼Ub was superposed onto Ube2W. **(D)** Close-up of the RING:Ube2W∼Ub model showing a potential salt bridge between TRIM21 E13 and ubiquitin K11. **(E)** Schematic model of the catalytic RING topology for N^α^-ubiquitination of TRIM21 by a Ube2W dimer. **(F)** Ube2W-mediated TRIM21 RING mono-ubiquitination assay using 10 µM T21-R-PS or R-R-PS and 0.25 µM Ube2W WT or monomeric V30K/D67K. Representative example of n = 2 independent experiments. **(G)** Schematic of antibody-induced recruitment of either two RINGs or two RING dimers. Only the latter satisfies the ‘two-plus-one’ model for RING autoubiquitination ^19,38^. **(H)** Antibody induced N^α^-ubiquitination of 100 nM R-PS or R-R-PS in the absence or presence of 1 molar equivalent of anti-GFP antibody. Ube2W was titrated (25, 50, 100, 200 nM). Representative example from n = 3 independent experiments. (**I)** Quantification of monoubiquitination from (H). Graph shows mean and s.e.m. from n = 3 independent experiments. **(J)** RPE-1 CAV1-mEGFP cells were electroporated with PBS or anti-GFP antibody ± indicated E2 proteins and 24h later CAV1-mEGFP fluorescence was quantified using the IncuCyte system. Values normalized to PBS control condition. Graph shows mean and s.e.m from n = 3 independent experiments (black dots). Statistical significance in (I) and (J) is based on two-tailed Student’s *t*-test.

### A Ube2W dimer monoubiquitinates TRIM21 RING and is required for Trim-Away

We postulated that as Ube2W is normally dimeric^46^, it may utilise a similar catalytic RING topology to that previously described for the Ube2N/Ube2V2 heterodimer^38^. Under such an arrangement, two RINGs could form a dimer to act as the enzyme, activating the donor ubiquitin on one Ube2W monomer, while a third RING acts as the substrate, oriented by the second Ube2W monomer to allow attack on the N-terminus (Figure 1E). We tested this hypothesis using two TRIM21 constructs, carrying either one (R) or two (R-R) RINGs and a PRYSPRY (PS) domain (R-PS and R-R-PS). Both constructs were efficiently monoubiquitinated by Ube2W, however this activity was abolished when monomeric Ube2W^V30K/D67K^ was used (Figure 1F). This is consistent with Ube2W dimerization being used to orient one RING domain as a substrate (Figure 1E). Surprisingly, no difference between R-PS and R-R-PS was observed, probably because the relatively high TRIM21 concentrations were sufficient to drive RING dimerization of R-PS. Previously we have shown that TRIM21 is activated in cells by substrate-induced clustering^15^ and indeed the addition of IgG Fc to in vitro ubiquitination experiments is required to induce TRIM21-mediated K63-chain formation by Ube2N/Ube2V2 under near-physiological enzyme concentrations^38^. We therefore reduced R-PS or R-R-PS concentrations and titrated Ube2W either in the presence or absence of IgG. Under these conditions, monoubiquitination was only observed with R-R-PS (Figure 1G&H). Moreover, R-R-PS monoubiquitination was greatly stimulated by the addition of IgG (Figure 1H&I). Importantly, we observed that endogenous TRIM21 is similarly dependent upon dimeric Ube2W for substrate degradation in cells. Monomeric Ube2W^V30K/D67K^, but not wild-type Ube2W, prevented degradation of substrate during Trim-Away (Figure 1J). A similar effect was observed with catalytically-inactive Ube2N (Ube2N^N119A^/Ube2V2)^38^ (Figure 1J). These results confirm previous observations using RNAi that both Ube2W and Ube2N/Ube2V2 are required for TRIM21 activity^28,29^. Taken together, the cellular and in vitro data suggest that antibody binding promotes TRIM21 RING monoubiquitination by dimeric Ube2W through a similar *trans* mechanism to Ube2N/Ube2V2 and that this is required for substrate degradation.

### Biochemical inhibition of TRIM21 N-terminal ubiquitination

While Ube2W is needed for substrate degradation and TRIM21 can be monoubiquitinated by the E2 in vitro, this does not prove that one requires the other. To investigate the requirement for TRIM21 N-terminal monoubiquitination, we decided to block it biochemically via N-terminal (N^α^)-acetylation. N^α^-acetylation is an irreversible modification catalysed in cells by N-Acetyl Transferases (NATs) using the co-factor Acetyl-CoA^47^. We chose Naa50, a NAT from *Chaetomium thermophilum* as its substrate specificity (MASS, MVNA) nicely matches the N-terminus of TRIM21 (MASA), and it displays thermostability and high *in vitro* activity^48^. NAT was incubated with R-R-PS in the presence of Acetyl-CoA. Successful N-terminal acetylation was confirmed by LC-MS/MS (Supplementary Figure 2A). N-terminally acetylated R-R-PS (Ac-R-R-PS) was added together with Ube2W and ubiquitin to test whether monoubiquitination was inhibited (Figure 2A). In the absence of acetylation, all R-R-PS was monoubiquitinated, whereas R-R-PS incubated with NAT and Acetyl-CoA remained predominantly non-ubiquitinated (Figure 2B). The degree of monoubiquitination inhibition was proportional to the time of NAT and Acetyl-CoA incubation (Supplementary Figure 2B). Importantly, the formation of free K63-linked ubiquitin chains was not compromised, demonstrating that the acetylated RING remains catalytically active (Supplementary Figure 2C). These results confirm both that TRIM21 RING is monoubiquitinated by Ube2W at its N-terminus and that this can be inhibited by N-terminal acetylation.

**Figure 2:**
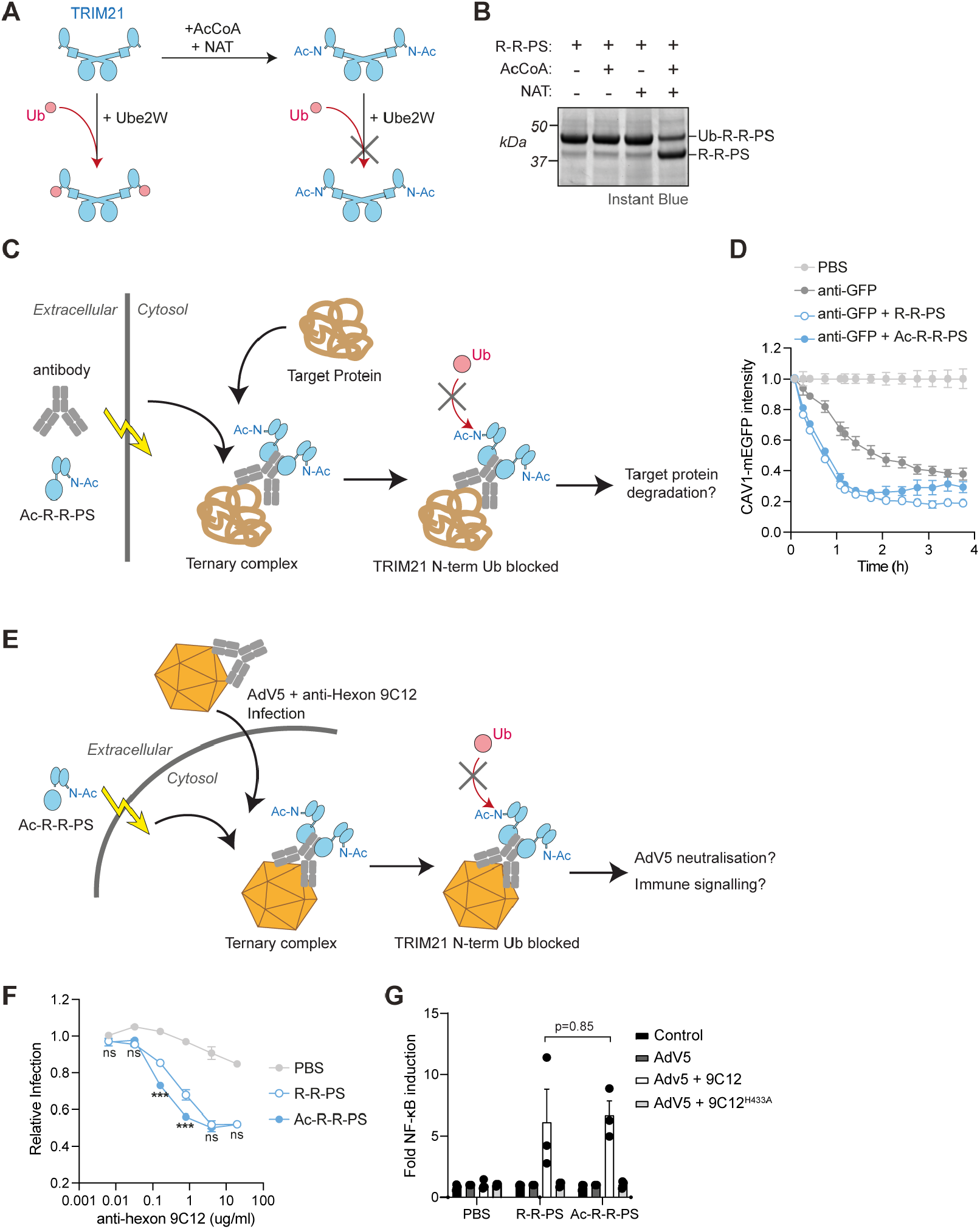
Ligase N-terminal ubiquitination is not required for substrate degradation and antiviral activity. **(A)** Schematic showing N-terminal acetylation of TRIM21 by AcCoA and NAT and then incubation of the acetylated or unmodified ligase with ubiquitin (Ub) and Ube2W in a ubiquitination reaction. **(B)** Protein-stained gel of ubiquitination reaction depicted in (A) using TRIM21 R-R-PS ligase. Monoubiquitination of ligase that has been incubated with AcCoA and NAT is inhibited. Representative example from n = 3 independent experiments. **(C)** Schematic showing Trim-Away experiment in which antibody is electroporated into cells together with R-R-PS (± N^α^-acetylation). Once inside cells, a ternary complex with the target protein is formed. If degradation is driven by ligase N-terminal ubiquitination, then N-terminally acetylated R-R-PS activity will be inhibited. **(D)** Results of Trim-Away experiment described in (C) shows that N-terminal acetylation of the ligase does not alter the kinetics of substrate (CAV1-GFP) degradation. RPE-1 CAV1-mEGFP cells were electroporated with PBS or anti-GFP antibody ± R-R-PS proteins and CAV1-mEGFP fluorescence was quantified using the IncuCyte system. Time shows hours (h) post-electroporation. Values normalized to PBS control condition. Graphs shows mean and s.e.m. from n = 4 technical replicates. Representative example from n = 2 independent experiments. Note that there is CAV1-mEGFP degradation with anti-GFP alone due to the presence of endogenous cellular TRIM21. **(E)** Schematic showing electroporation of N^α^-acetylated ligase (Ac-R-R-PS) into cells followed by infection with Adv5 in the presence of anti-hexon antibody 9C12. If ligase N-terminal ubiquitination is necessary for TRIM21 antiviral function, neutralization of infection and immune signaling will be inhibited. **(F**,**G)** Neutralization of AdV5 infection by increasing 9C12 concentrations (F) and AdV5-9C12-induced NFkB activation (G) in HEK293T TRIM21 KO cells infected immediately after electroporation with PBS or R-R-PS ± N^α^-acetylation. 9C12^H433A^ does not bind TRIM21 PRYSPRY. Graphs shows mean and s.e.m. from n = 3 independent experiments. Black dots in (G) show individual data points. Statistical significance between R-R-PS and Ac-R-R-PS is based on two-way ANOVA (F) and Student’s *t*-test (G).

### TRIM21 N-terminal ubiquitination is not required for its activity

With a method to specifically block N-terminal ubiquitination of the RING, we tested whether this is required for substrate degradation. Either R-R-PS or Ac-R-R-PS were electroporated together with anti-GFP antibody into cells expressing the substrate CAV1-mEGFP and the kinetics of degradation monitored over time detection (Figure 2C). Consistent with previous Trim-Away experiments, electroporation of anti-GFP antibody drove substrate degradation via endogenous TRIM21 (Figure 2D). However, CAV1-mEGFP degradation was substantially accelerated through the delivery of exogenous R-R-PS (Figure 2D). Importantly, ligase acetylation (Ac-R-R-PS) had no impact and degradation proceeded with identical kinetics (Figure 2D). Next, we tested whether N-terminal acetylation was required for TRIM21 antiviral function. Cells were electroporated with either R-R-PS or Ac-R-R-PS, then infected with Adenovirus 5 (AdV5) in the presence of the anti-hexon antibody 9C12 (Figure 2E). Previous experiments have shown that when TRIM21 binds to antibody-coated virus it blocks infection by mediating proteasomal-degradation of the virion^2^. As expected, electroporation of R-R-PS neutralized AdV5 in an antibody-dose dependent manner (Figure 2F). Ac-R-R-PS was at least as active as R-R-PS at neutralizing AdV5 infection and at intermediate antibody concentrations was more effective (Figure 2F). In addition to mediating virion degradation, TRIM21 also activates innate immune signalling upon detection of antibody-coated virus^29^. Both R-R-PS and Ac-R-R-PS were equally effective at stimulating NF-*κ*B-driven transcription in response to antibody-coated Adv5 (Figure 2G). This was dependent upon direct antibody engagement as the 9C12 mutant H433A, which specifically ablates binding to the PRYSPRY^20^, failed to activate NF-*κ*B. Taken together, the data show that N-terminal ubiquitination of the RING ligase is dispensable for both Trim-Away and TRIM21 antiviral functions.

### TRIM21 N-terminal ubiquitination regulates its own stability

We reasoned that if RING ligase autoubiquitination is not required for substrate degradation, perhaps it is involved in mediating self-turnover. We therefore electroporated Ac-R-R-PS into cells and monitored protein levels after 1 hour (Figure 3A). Comparison of epoxomicin treated and untreated cells revealed that non-acetylated R-R-PS was readily degraded by the proteasome (Figure 3B, first two lanes). In contrast, epoxomicin had little impact on Ac-R-R-PS protein levels, indicating that blocking ligase N-terminal ubiquitination prevents its proteasomal degradation (Figure 3B, last two lanes). This data suggests that while ligase N-terminal ubiquitination is not required for substrate degradation it may be used to regulate ligase levels. To test this, we repeated our electroporation experiments with R-R-PS or Ac-R-R-PS but measured either Trim-Away or AdV5 neutralization after a delay of several hours (Figure 3C). For Trim-Away experiments, this was accomplished by co-electroporating the RNA encoding the antibody construct responsible for recruiting R-R-PS to substrate (vhhGFP4-Fc). Trim-Away is thus delayed by several hours while vhhGFP4-Fc is expressed. Under this experimental regime, there was no longer any substrate degradation in cells electroporated with R-R-PS (Figure 3D, compare with Figure 2D). In contrast, Ac-R-R-PS degraded CAV1-mEGFP just as efficiently as when Trim-Away proceeds immediately upon ligase electroporation (Figure 3D). For AdV5 neutralization experiments, cells were infected 2 hours post-ligase delivery. In this case, neutralization by R-R-PS neutralization was severely attenuated with Ac-R-R-PS inhibiting infection significantly more efficiently (Figure 3E).

**Figure 3:**
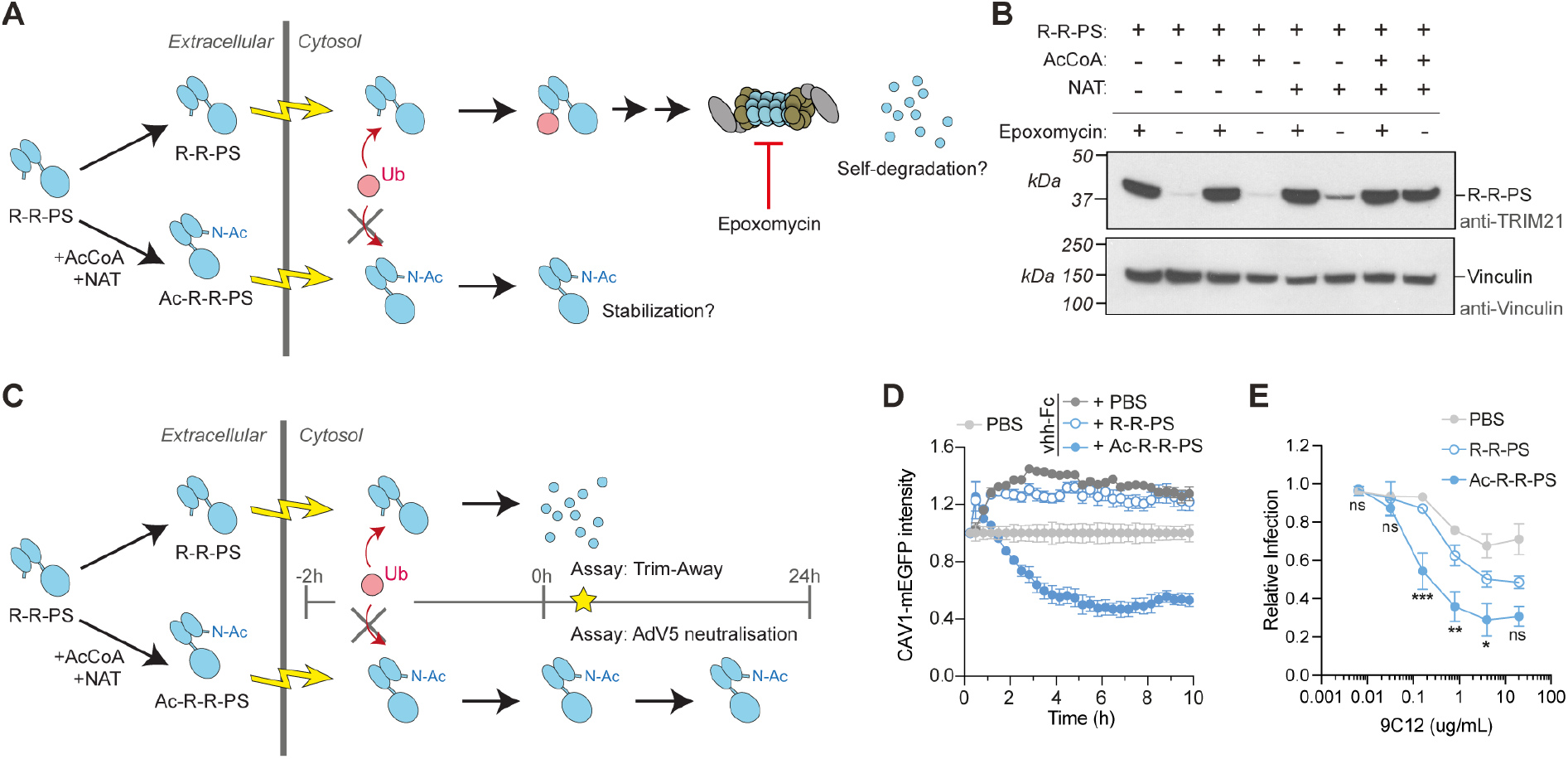
N-terminal ubiquitination regulates ligase turnover in cells. **(A)** Schematic showing electroporation of R-R-PS ± N^α^-acetylation and ubiquitin-proteasome dependent self-degradation. **(B)** Western blot of experiment depicted in (A) 1 hour post-electroporation. R-R-PS protein levels are rescued by addition of 10 µM proteasome inhibitor epoxomycin. Acetylated R-R-PS protein persists in cells irrespective of proteasome inhibition. Representative example from n = 2 independent experiments. **(C)** Schematic showing electroporation of R-R-PS ± N^α^-acetylation into cells, followed by delayed Trim-Away or Adv5 neutralization assays. **(D)** For the delayed Trim-Away assay, mRNA encoding the antibody construct (vhhGFP4-Fc) responsible for recruiting R-R-PS to substrate (CAV1-mEGFP) is co-electroporated into NIH3T3-CAV1-mEGFP cells with PBS or R-R-PS ± N^α^-acetylation; Trim-away is delayed for ∼2h until vhhGFP4-Fc protein is translated. Graph shows mean and s.e.m. from 4 technical replicates of CAV1-mEGFP fluorescence quantified using the IncuCyte system. Values normalized to PBS control condition (no vhhGFP4-Fc). Time shows hours (h) post-electroporation Representative example from n = 2 independent experiments. Note that NIH3T3 cells do not contain endogenous TRIM21 and expression of vhhGFP4-Fc in the absence of TRIM21 activity leads to GFP stabilization. **(E)** For the delayed Adv5 neutralization assay, HEK293T TRIM21 KO cells are infected with AdV5 ± 9C12 two hours post-electroporation of PBS or R-R-PS ± N^α^-acetylation. Graph shows mean and s.e.m. from n = 3 independent experiments. Statistical significance between R-R-PS and Ac-R-R-PS is based on two-way ANOVA.

### TRIM21 ubiquitinates antibody:substrate independently from itself

As ligase autoubiquitination is not required for substrate degradation, we investigated whether this might be driven by substrate ubiquitination instead. To test this, we performed in vitro ubiquitination experiments with R-R-PS, anti-GFP antibody and recombinant mEGFP substrate (Figure 4A). We observed simultaneous ubiquitination of R-R-PS, antibody heavy chain and mEGFP (Figure 4B). Monoubiquitination was observed upon incubation with Ube2W alone, whereas in conditions where Ube2N/V2 was also included this resulted in anchored polyubiquitin chains (Figure 4B). Importantly, substrate ubiquitination only occurred when both R-R-PS and antibody were present (Supplementary Figure 3). This is consistent with the requirement for antibody to recruit the TRIM21 ligase to its substrate. Next, we asked whether substrate ubiquitination occurs independently of ligase ubiquitination or if the latter modification is required to stimulate catalytic activity. To do this, we repeated our in vitro ubiquitination assay in the presence of both E2s and compared R-R-PS with Ac-R-R-PS. Ligase acetylation blocked autoubiquitination but did not interfere with polyubiquitination of either antibody heavy chain or substrate (Figure 4C). This data shows that TRIM21 catalyses polyubiquitination of an antibody-bound substrate and that this can occur independently of ligase autoubiquitination. Nevertheless, the fact that TRIM21, antibody and substrate can be simultaneously polyubiquitinated in vitro is consistent with in-cell Trim-Away data showing that all three components are simultaneously degraded^24^.

**Figure 4:**
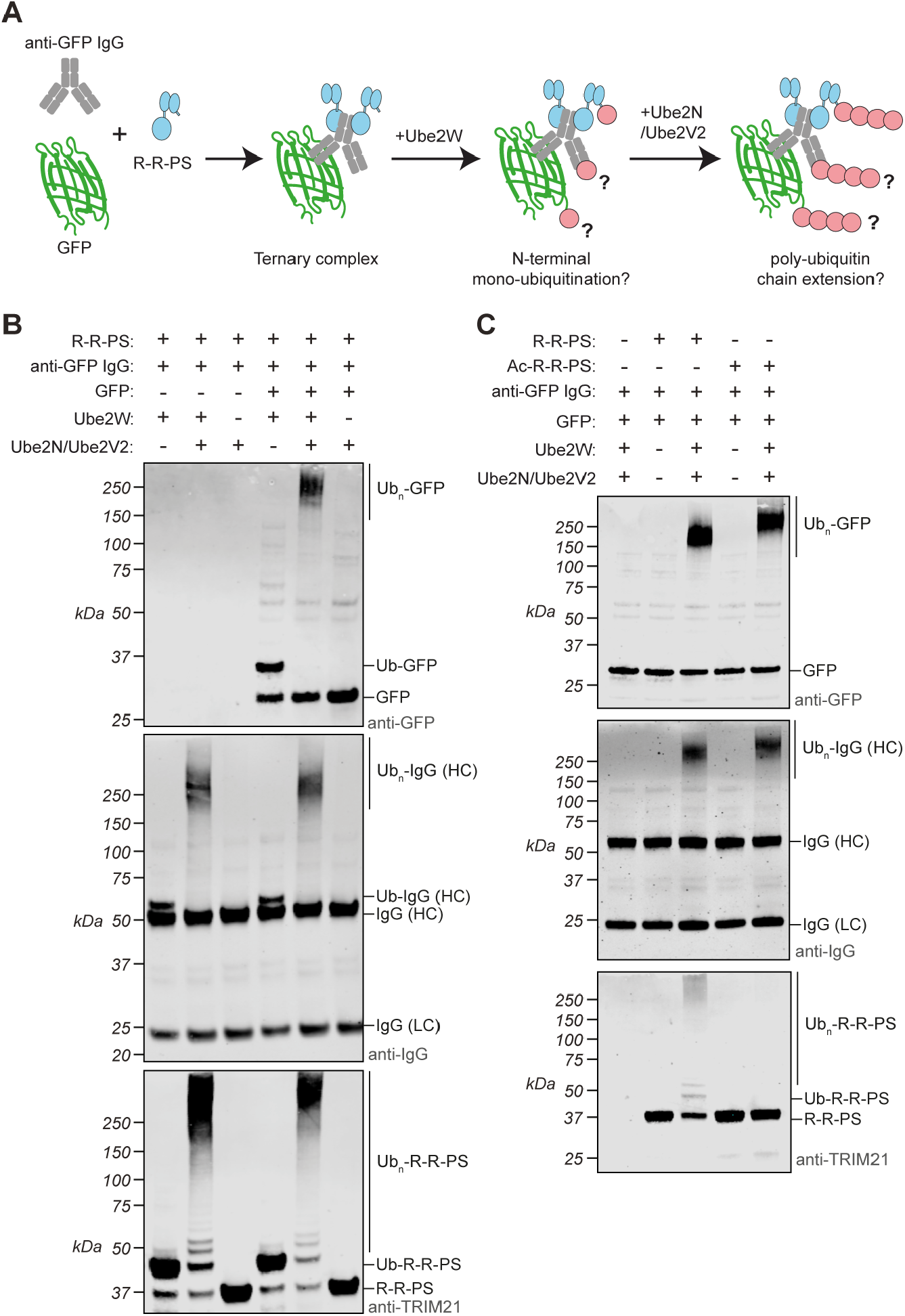
TRIM21 independently ubiquitinates itself and its substrate. **(A)** Schematic showing an in vitro ubiquitination reaction in which ligase (R-R-PS), substrate (GFP) and anti-GFP antibody are incubated together with various E2 enzymes to promote either mono- or polyubiquitination. **(B)** Western blot of experiment described in (A). Top panel is blotted for GFP, middle panel for IgG and lower panel for TRIM21. Depending on the E2s present, monoubiquitinated species or a higher molecular weight smear indicative of polyubiquitin is observed. Representative example from n = 2 independent experiments. **(C)** Western blot of experiment similar to (B) but comparing R-R-PS to Ac-R-R-PS. Note that while R-R-PS ubiquitinates itself, antibody heavy chain and substrate, Ac-R-R-PS only ubiquitinates antibody and substrate. Representative example from n = 2 independent performed experiments. See also Supplementary Figure 3.

### Substrates are ubiquitinated during Trim-Away in live cells

Next, we attempted to monitor substrate ubiquitination during Trim-Away degradation in living cells (Figure 5A). We chose two substrates, ERK1 and IKK*α*, and blotted for protein levels at various timepoints post antibody electroporation. For both substrates, high-molecular weight bands or smearing consistent with polyubiquitination was observed at 30 minutes post-antibody electroporation (Figure 5B&C). These high molecular weight bands decreased over the next few hours simultaneous with a reduction in substrate protein levels. To obtain further evidence for substrate polyubiquitination we repeated the IKK*α* Trim-Away experiment in the presence of proteasome inhibitor MG132. Addition of MG132 had no impact in control cells, but in the presence of electroporated antibody higher molecular weight laddering was clearly observed (Figure 5D). When performed as a time-course experiment, this revealed that IKK*α* ubiquitinated species first formed and then was depleted coincident with protein degradation. Treatment with MG132 both blocked degradation and led to a steady accumulation of ubiquitinated material (Supplementary Figure 4A). We also probed for TRIM21 and observed a decrease in protein levels that also paralleled substrate depletion (Figure 5B&C). Higher molecular weight bands were observed for TRIM21 that may also indicate polyubiquitination, however whilst these increased during ERK1 Trim-Away there was no change with IKK*α*. To test whether antibody-dependent substrate ubiquitination is mediated by TRIM21 we repeated our experiments in TRIM21 knockout (TRIM21 KO) cells reconstituted with either HA-tagged TRIM21 (+T21-HA) or an empty vector (+EV) control. Antibody-induced ERK1 laddering indicative of polyubiquitination was observed in knockout cells reconstituted with TRIM21-HA but not empty vector (Figure 5E). These data indicate that substrates are ubiquitinated during Trim-Away in live cells in an antibody- and TRIM21-dependent manner.

**Figure 5:**
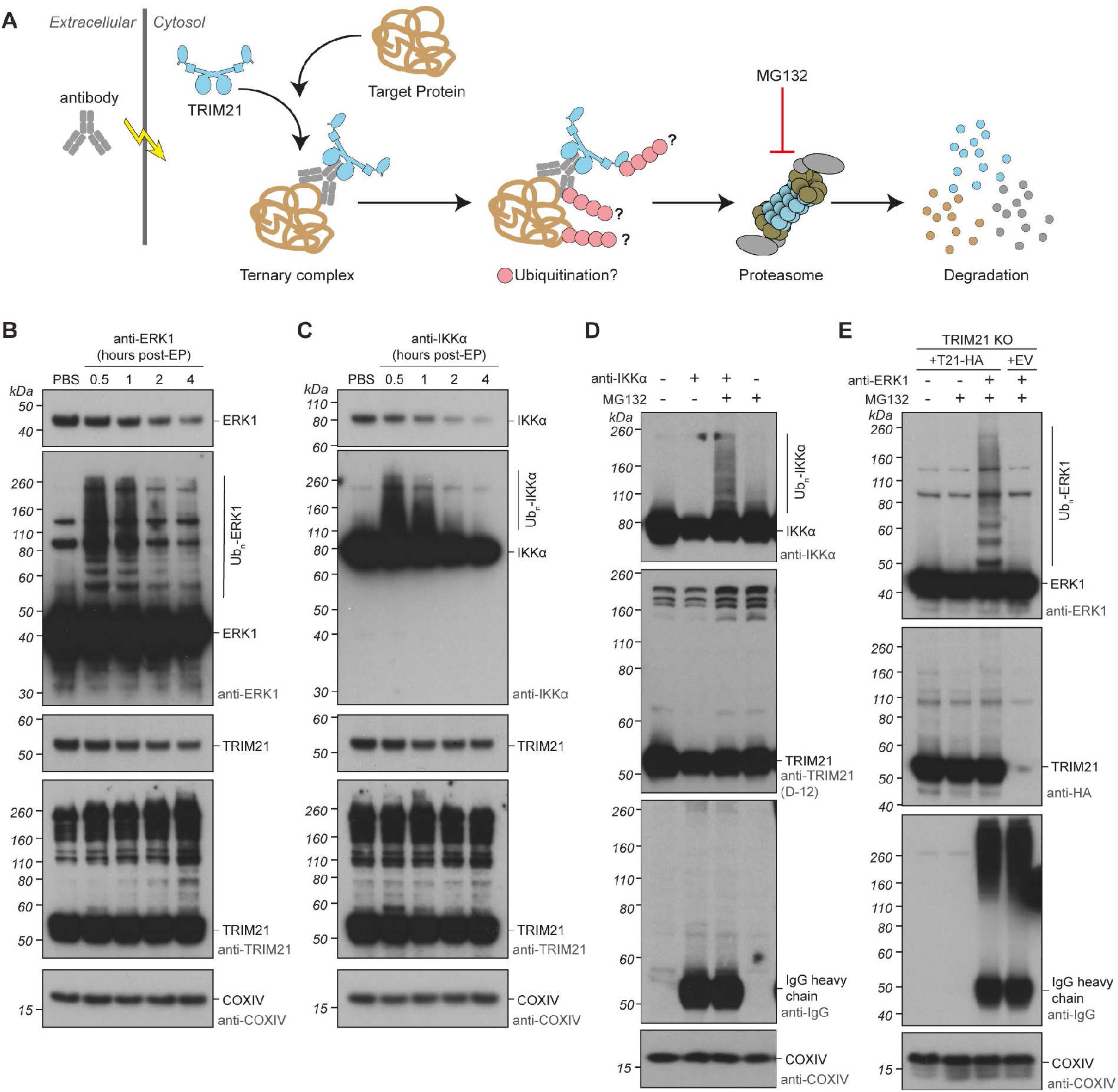
Substrate ubiquitination parallels substrate degradation during Trim-Away. **(A)** Schematic showing Trim-Away experiment in which antibodies are electroporated into cells expressing endogenous TRIM21. Ubiquitination and degradation are then monitored in the presence or absence of proteasome inhibitor MG132. **(B**,**C)** RPE-1 cells were electroporated with PBS or (B) anti-ERK1 antibody or (C) anti-IKKα antibody and whole cell lysates harvested at the indicated times after electroporation for immunoblotting. Short exposures show degradation of TRIM21 and substrates. Long exposures reveal substrate ubiquitination followed by degradation of ubiquitinated species. **(D)** RPE-1 cells were electroporated with PBS or anti-IKKα antibody ± MG132 and whole cell lysates harvested 3h post-electroporation for immunoblotting. **(E)** RPE-1 TRIM21 KO cells reconstituted with TRIM21-HA or empty vector (EV) were electroporated with PBS or anti-ERK1 antibody ± MG132 and whole cell lysates harvested 1h post-electroporation for immunoblotting. Representative examples from 3 independent experiments.

Previously, we have shown that Trim-Away can be performed in the absence of antibody by fusing a substrate-targeting nanobody directly to domains from TRIM21^15^. We used this approach to determine whether there is something particular to the ternary complex formed between TRIM21:antibody:substrate that is required for substrate ubiquitination. We tested two fusion constructs in which either the TRIM21 RING, B Box and coiled-coil (T21RBCC-) or RING domain alone (T21R-) is fused to the anti-GFP nanobody (vhhGFP4). Efficient Trim-Away was observed using both fusion constructs (Supplementary Figure 4B). Treatment with MG132 inhibited degradation and led to a coincident accumulation of ubiquitinated substrate (Supplementary Figure 4B). These results show that neither antibody nor a ternary complex is required for TRIM21-mediated substrate ubiquitination and degradation. Introducing two RING-inactivating mutations into the T21R-vhhGFP4 construct (T21R^I18R/M72E^-vhhGFP4)^15^ completely abolished both substrate ubiquitination and degradation (Supplementary Figure 4C) suggesting these processes are driven by TRIM21 RING catalytic activity.

### Neither N-terminal nor lysine ubiquitination of TRIM21 is required for Trim-Away

The preceding data show that activated TRIM21 can catalyse simultaneous ubiquitination of itself, antibody and substrate. However, blocking N-terminal TRIM21 ubiquitination does not prevent substrate degradation. To exclude the possibility that ligase lysine autoubiquitination drives substrate degradation, we made a variant of T21R-vhhGFP4 in which all lysines were substituted for arginines (Figure 6A; T21R^K0^-vhhGFP4^K0^). Remarkably, removal of all lysines from T21R-vhhGFP4 had no impact on either the kinetics or efficiency of substrate degradation (Figure 6B&C). Furthermore, simultaneous blocking the N-terminus via acetylation also had no effect (Figure 6B&C). Taken together, these data show that substrate degradation by TRIM21 is not dependent upon ligase canonical autoubiquitination. Importantly, however, substrate turnover can be uncoupled from ligase turnover as inhibiting N-terminal autoubiquitination reduced ligase depletion without altering substrate degradation (Figure 6C).

**Figure 6:**
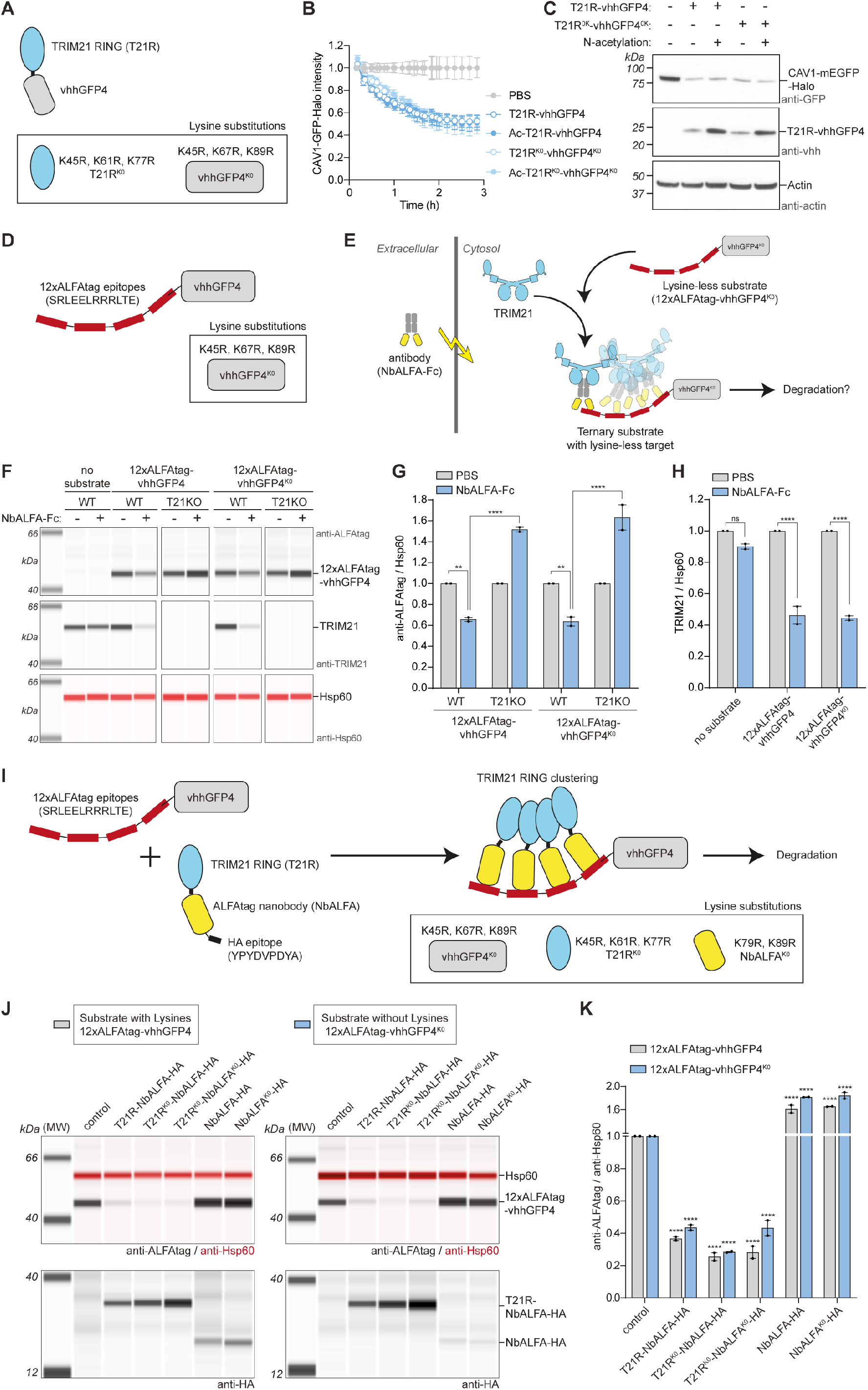
Trim-Away mediates protein depletion in the absence of lysine ubiquitination on either ligase or substrate. **(A)**. Schematic showing T21R-vhhGFP4 fusion protein and substitutions to remove all lysines. **(B**,**C)** RPE-1 CAV1-mEGFP-Halo cells were electroporated with PBS or T21R-vhhGFP4 protein ± lysines ± N^α^-acetylation. **(B)** CAV1-mEGFP-Halo fluorescence was quantified using the IncuCyte system. Time shows hours (h) post-electroporation. Values normalized to PBS control condition. Graphs shows mean and s.e.m. from n = 4 technical replicates. **(C)** Whole cell lysates were harvested 3h post-electroporation for immunoblotting. Representative examples (B,C) from 3 independent experiments. **(D)** Schematic showing a model lysine-less substrate consisting of a vhhGFP4 nanobody (lysine substitutions in box) with 12 copies of the naturally lysine-less ALFAtag epitope. **(E)** Scheme showing Trim-Away experiment in which the anti-ALFAtag antibody (NbALFA-Fc) is electroporated into cells expressing endogenous TRIM21 and the lysine-less substrate (12xALFAtag-vhhGFP4^K0^). **(F-H)** RPE-1 WT or TRIM21 KO cells expressing either substrate with lysines (12xALFAtag-vhhGFP4) or without lysines (12xALFAtag-vhhGFP4^K0^) were electroporated with PBS or NbALFA-Fc and whole cell lysates harvested 8h post-electroporation for capillary-based immunoblotting. Lane view (F) and quantification (G,H) of substrate (G) and TRIM21 (H) protein levels normalized to PBS condition. Graphs show mean and s.e.m. from 2 independent experiments (black dots). Statistical significance is based on two-way ANOVA. Note that binding of NbALFA-Fc in the absence of TRIM21 causes stabilization of substrate. **(I)** Schematic of completely lysine-less Trim-Away assay. The substrate constitutes a vhhGFP4 nanobody with 12 copies of the ALFAtag epitope. The ligase constitutes the T21 RING fused to an anti-ALFAtag nanobody. The 12xALFAtag allows clustering of multiple T21 RINGs, triggering degradation. The substitutions necessary to remove lysines from each domain in the assay are shown boxed. Note that the ALFAtag and HA epitopes are naturally lysine-less. **(J**,**K)** RPE-1 TRIM21 KO cells expressing either substrate with lysines (12xALFAtag-vhhGFP4) or without lysines (12xALFAtag-vhhGFP4^K0^) were electroporated with water (control) or mRNA encoding the indicated constructs and whole cell lysates harvested 8h post-electroporation for capillary-based immunoblotting. Lane view (J) and quantification (K) of substrate protein levels normalized to control condition. Graph shows mean and s.e.m. from 2 independent experiments (black dots). Statistical significance from control condition is based on two-way ANOVA. Note that binding of NbALFA alone causes stabilization of substrate.

### Trim-Away degrades lysine-less substrates

To test whether Trim-Away is driven by substrate lysine ubiquitination, we designed model substrates comprising a dodecameric ALFAtag repeat, which naturally contains no lysine residues^49^, fused to vhhGFP4 with or without lysines (Figure 6D). These substrates were expressed in either wild-type (WT) or TRIM21 knockout (T21KO) cells, which were then electroporated with an anti-ALFAtag antibody (vhhNbALFA-Fc). The NbALFA-Fc is predicted to bind to the ALFAtag substrate and recruit endogenous TRIM21 via Fc interaction, leading to TRIM21 clustering, activation and degradation (Figure 6E). Indeed, NbALFA-Fc electroporation triggered substrate degradation in WT but not T21KO cells (Figure 6F&G). Importantly, degradation was not dependent upon substrate lysines, as substrates both with and without lysines were equally well degraded (Figure 6F&G). As expected for a Trim-Away experiment, endogenous TRIM21 was also degraded alongside each substrate (Figure 6H). As it is formally possible that degradation of a TRIM21:substrate complex requires only one partner to undergo lysine ubiquitination, we modified our assay to remove lysines simultaneously from both ligase and substrate. To do this, we complemented our model substrate with a model ligase comprising a TRIM21 RING fused to the anti-ALFAtag nanobody^49^ (T21R-NbALFA; Figure 6I). In this assay, the ALFAtag substrate is predicted to recruit multiple T21R-vhhALFA ligases, leading to ligase clustering, activation and degradation (Figure 6I). As before, removal of all lysines from the substrate had no impact on substrate degradation (Figure 6J&K). Strikingly however, Trim-Away was equally efficient when both ligase and substrate were lysine-less (Figure 6J&K). Taken together, our data show that ligase autoubiquitination does not drive TRIM21-mediated degradation but neither does substrate lysine ubiquitination. This finding may explain the efficiency with which Trim-Away degrades diverse substrates^24^.

## DISCUSSION

Existing models of TRIM ligase ubiquitination for antivirals TRIM5 and TRIM21 propose that N-terminal RING autoubiquitination drives substrate degradation^26,50^ (Supplementary Figure 5). These models are based on data showing that substrate and ligase are co-degraded^22^, with identical kinetics^15,24^, and that while TRIM autoubiquitination is readily detected in cells^15,25,28^, ubiquitination of viral substrates is not. Additionally, depletion of the E2 responsible for N-terminal ubiquitination, Ube2W, inhibits TRIM5 and TRIM21 antiviral functions^26,28^. Here we show that while TRIM21 N-terminal autoubiquitination via Ube2W is stimulated by substrate binding and drives turnover of the ligase inside cells, it is not required for substrate degradation. TRIM21 directly catalyses ubiquitination of its substrates in vitro and in cells and this proceeds efficiently even when ligase N-terminal ubiquitination is prevented (Supplementary Figure 5). The ubiquitination of ligase lysine residues is also unnecessary, as a lysine-less and N-terminally acetylated RING readily degrades substrates. Furthermore, we find that TRIM21 carries out efficient substrate degradation even when both the ligase and substrate lack lysines. These unexpected findings pose two important questions for TRIM ligase mechanism, namely what modification drives substrate degradation and why does the ligase ubiquitinate and degrade itself?

The answer to this last question may lie in the differences between TRIM and CRL ligases. CRLs are assembled combinatorially from a small number of RINGs, adaptors and scaffolds^11,12^. Degradation of CRL components would be both detrimental and unnecessary. In the absence of substrate, CRL complexes rapidly disassemble, providing an intrinsic functional shut-off^51^. In contrast, TRIM ligases encode catalytic and substrate binding domains in one polyprotein and so cannot be regulated by recruiting independent ligase components. Instead, TRIM proteins appear to be activated by forming higher-order structures^1,15,39^. The ability to form large oligomeric structures, sometimes called ‘cytoplasmic bodies’, is a common feature of TRIM proteins^52^. Cytoplasmic bodies can be induced simply by over-expression, likely because TRIMs have evolved to readily assemble into large scaffolds. These scaffolds activate TRIM RINGs by mediating dimerization and amplifying the ubiquitination signal associated with the ligase:substrate complex^15,19^. Evolving a ubiquitination mechanism whereby the ligase is ubiquitinated and co-degraded alongside its substrate may be a necessary step to ensure that, once-formed, large TRIM assemblies do not persist as cytoplasmic bodies inside the cell, continuously catalysing ubiquitination. This may be particularly important for TRIMs like TRIM5 and TRIM21, which use their ubiquitin chains not only for degradation but also to potently stimulate inflammatory signalling^27,29^.

A regulatory role for ligase ubiquitination and degradation is further suggested by uncoupling substrate ubiquitination and degradation that we have demonstrated here. Previously, we have shown how a ‘two-plus-one’ catalytic RING topology drives TRIM autoubiquitination^19,28,38^. What drives substrate ubiquitination is currently unclear but the answer in this case may involve the quaternary structure unique to TRIM proteins. Despite their functional heterogeneity and inclusion of additional diverse domains, most TRIMs preserve not only the presence of a RING, B Box and coiled-coil but their relative arrangement and interdomain spacing^21^. The coiled-coils are antiparallel helices^53^ whose length is remarkably consistent across different TRIMs. This places each RING in a TRIM dimer at opposite ends of the molecule and a conserved distance apart. Many TRIMs also utilise a substrate-binding PRYSPRY domain and these are located centrally above the coiled-coil^54^. Given the fixed positions of these catalytic and substrate binding domains with respect to each other, it is tempting to speculate that TRIM ligases utilise a form evolutionary conserved scaffolds to ubiquitinate their substrates. In contrast to the CRL scaffold that is formed within the assembles CRL, TRIMs have evolved supramolecular dynamic scaffolds to perform the same biochemical function.

Perhaps the most surprising finding presented here is that lysines are not required for substrate degradation by TRIM21. One explanation may be that, like the ligase, substrates are ubiquitinated at their N-terminus. This would be consistent with the need for Ube2W during both TRIM5 and TRIM21 cellular function. Data shown here using dimerization mutant Ube2W^V30K/D67K^ supports previous siRNA depletion studies and the importance of this E2. Alternatively, degradation may involve the non-canonical ubiquitination of non-lysine residues. Several E3s are capable of ubiquitinating serine and threonine residues or non-protein substrates^55-58^. Whether lysine ubiquitination is redundant during the function of other diverse TRIM ligases remains to be explored. Nevertheless, the fact that TRIM21 is not reliant on substrate lysine availability may explain why Trim-Away technology is efficient at degrading diverse substrates.

## ACKNOWLEDGEMENTS

L.K. was supported by a Ph.D. Fellowship from the Boehringer Ingelheim Fonds. This work was supported by the MRC (UK; U105181010), a Wellcome Trust Investigator Award (200594/Z/16/Z) and a Wellcome Trust Collaborator Award (214344/A/18/Z).

## AUTHOR CONTRIBUTIONS

L.K., D.C. and L.C.J. conceptualized the study and prepared the manuscript; All authors developed methodology and/or performed experiments.

---

**Supplementary Figure 1:**
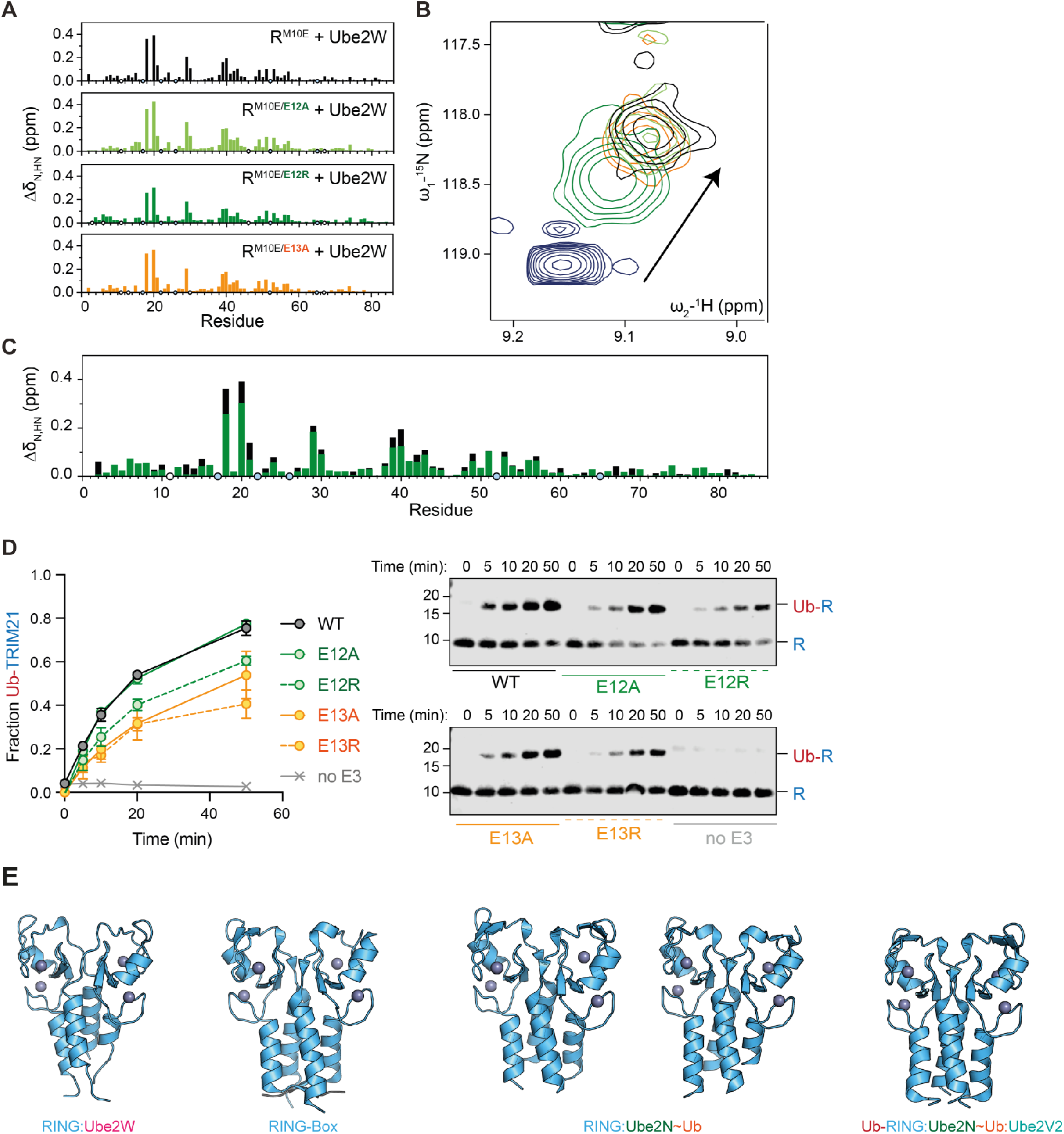
Interaction of Ube2W with TRIM21 RING. **(A)** Histograms of chemical shift perturbations (CSP) shown against the sequence of TRIM21 RING^M10E^ (R^M10E^). These CSPs result from NMR titrations of Ube2W^V30K/D67K/C91K^ against ^15^N-labelled TRIM21 tri-ionic mutants at a 1:1 molar ratio. Blue circles indicate proline residues, white circles missing assignments. **(B)** A part of ^15^N-HSQC spectral overlay of R^M10E^ in absence (blue) or presence of 1:1 molar equivalent of Ube2W^V30K/D67/C91K^. In addition, spectra of TRIM21 mutants (E12A in light green, E12R in dark green and E13A in orange) are shown in presence of 1:1 molar equivalent of Ube2W^V30K/D67/C91K^. **(C)** Histograms shown in (A) are here shown as an overlay. **(D)** Ube2W-mediated TRIM21 RING monoubiquitination assay. Shown is a time-course, where error bars represent s.e.m. from n = 3 independent experiments. Western blots are representative of all replicates. **(E)** Shown are RING dimers of different TRIM21 complexes (RING:Ube2W, RING-Box (5OLM)^14^, RING:Ube2N∼Ub (two RING dimers in asymmetric unit, 6S53)^30^, Ub-RING:Ube2N∼Ub:Ube2V2 (7BBD)^38^). Zn^2+^-atoms are shown as grey spheres, the isopeptide bond is marked by an arrow and polar interactions are indicated by dashed black lines.

**Supplementary Figure 2:**
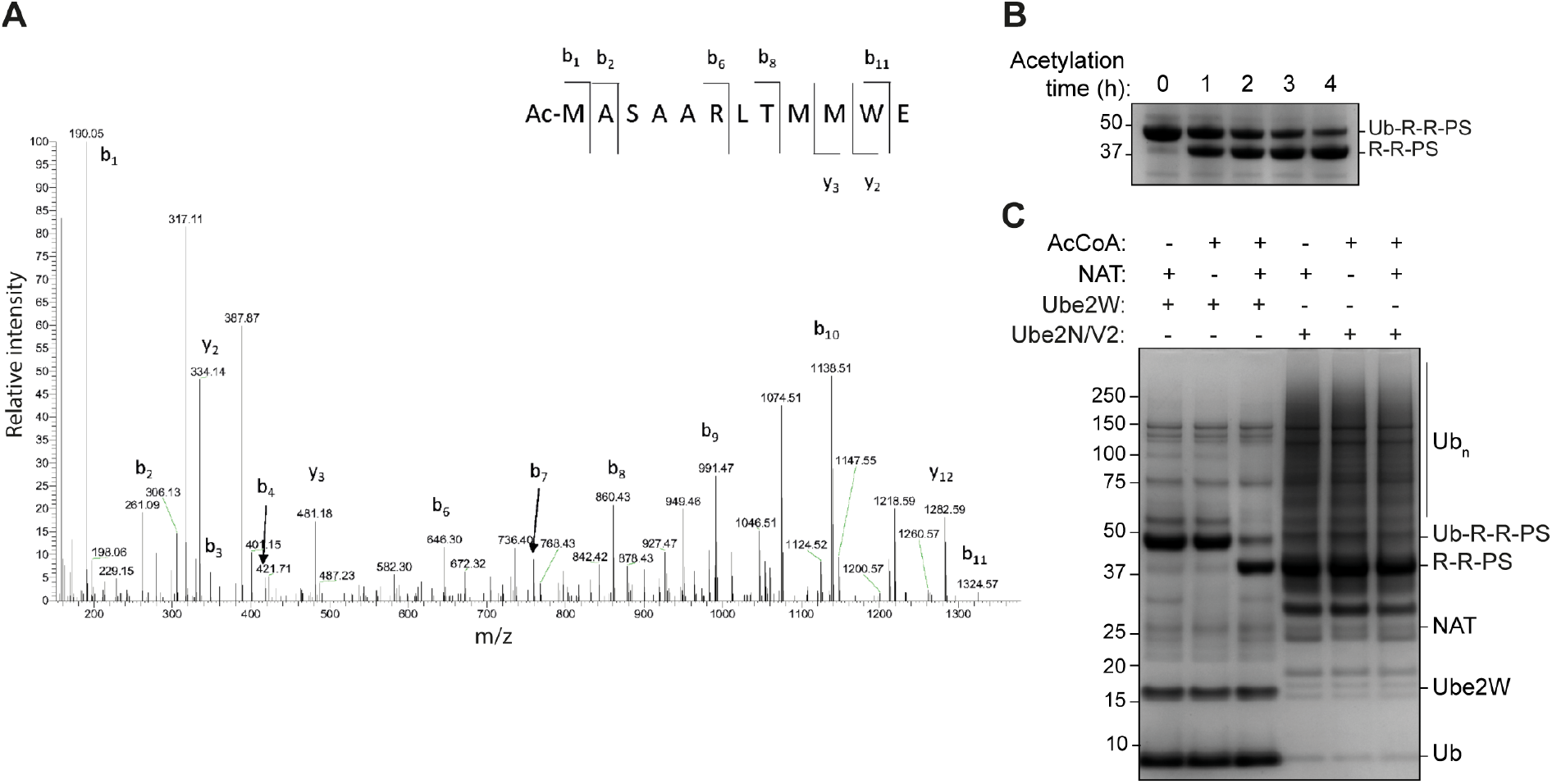
TRIM21 RING can be N-terminally acetylated with NAT and AcCoA to block N-terminal autoubiquitination. **(A)** LC-MS/MS spectra of TRIM21 R-R-PS after 4 h acetylation reaction show N^α^- acetylated TRIM21 N-terminal peptides after digestion with the protease N-Asp. **(B)** Instant-Blue-stained gels showing Ube2W-mediated TRIM21 mono-ubiquitination reactions with R-R-PS. Before the ubiquitination reaction, acetylation reactions were performed for the indicated times. Gel representative of n = 2 independent experiments. **(C)** As in (B) but showing results of ubiquitination reaction upon incubation with Ube2W and Ube2N/V2 for 1 h after 4 h of N^α^-acetylation. In contrast to N-terminal monoubiquitination, incubation with NAT and AcCoA doesn’t impact ubiquitin smearing characteristic of free K63-chain formation. Gel representative of n = 2 independent experiments.

**Supplementary Figure 3:**
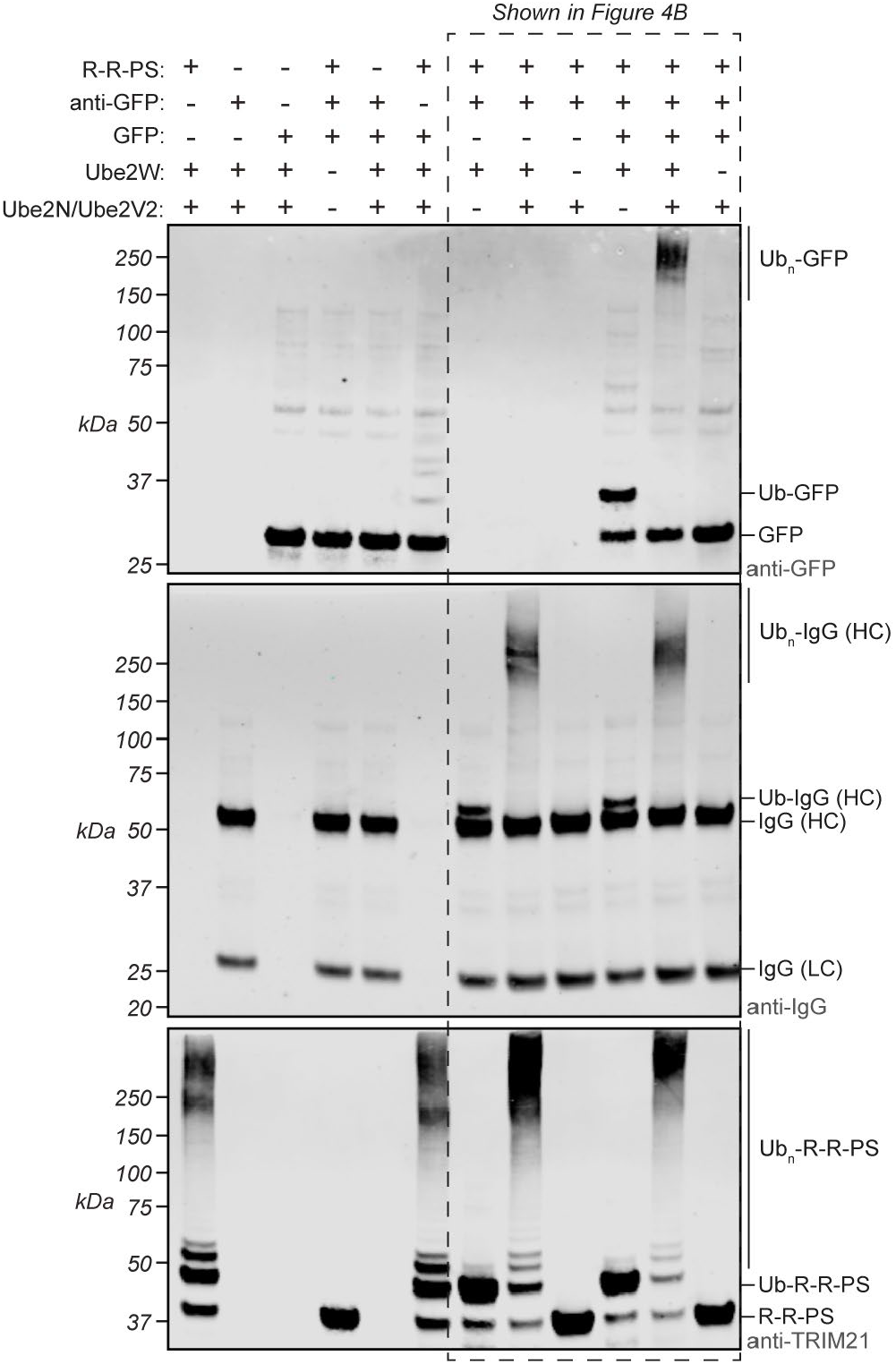
Ligase and substrate ubiquitination in vitro. Western blots of ubiquitination reactions using combinations of R-R-PS, anti-GFP antibody, GFP, Ube2W and Ube2N/Ube2V2. Substrate and antibody heavy chain are only ubiquitinated in the presence of R-R-PS and substrate only in the presence of both antibody and R-R-PS. Western blots representative of n = 2 independently performed experiments. See also Figure 4.

**Supplementary Figure 4:**
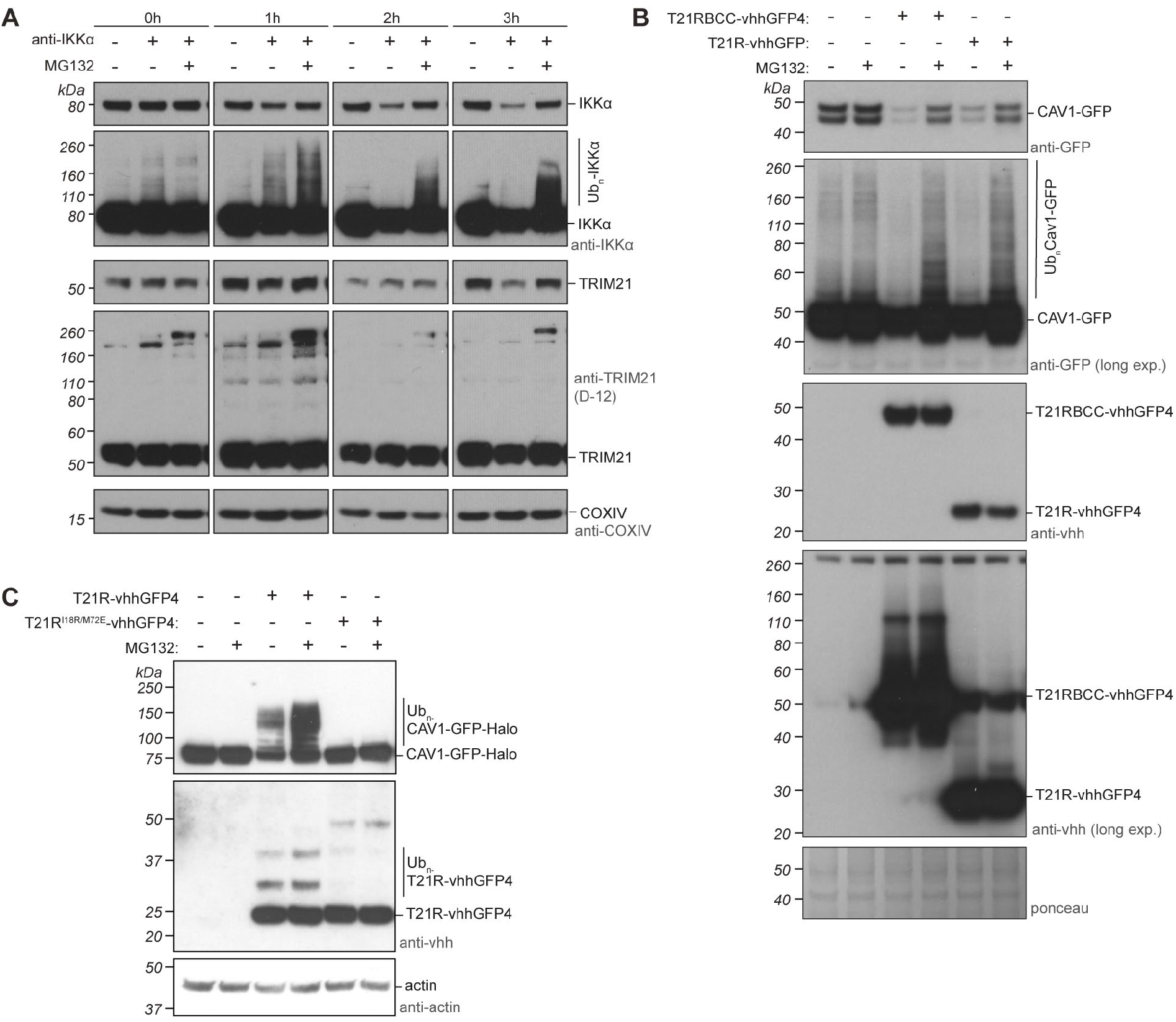
Substrate ubiquitination during Trim-Away in live cells. **(A)** Western blots of Trim-Away time-course experiment using anti-IKK*α* antibody in the presence or absence of MG132. Proteasome inhibition rescues IKK*α* degradation and leads to the accumulation of ubiquitinated protein. Western blots representative of n = 2 independent experiments. **(B)** Ubiquitination of CAV1-mEGFP by T21RBCC-vhhGFP4 or T21R-vhhGFP4 fusions in the presence or absence of MG132. Western blots representative of n = 2 independent experiments. Ponceau protein stain shows equal loading. **(C)** Western blot of Trim-Away experiment using WT T21R-vhhGFP4 or a mutant incapable of catalysing ubiquitination (T21R-vhhGFP4^I18R/M72E^). Western blots representative of n = 3 independent experiments.

**Supplementary Figure 5:**
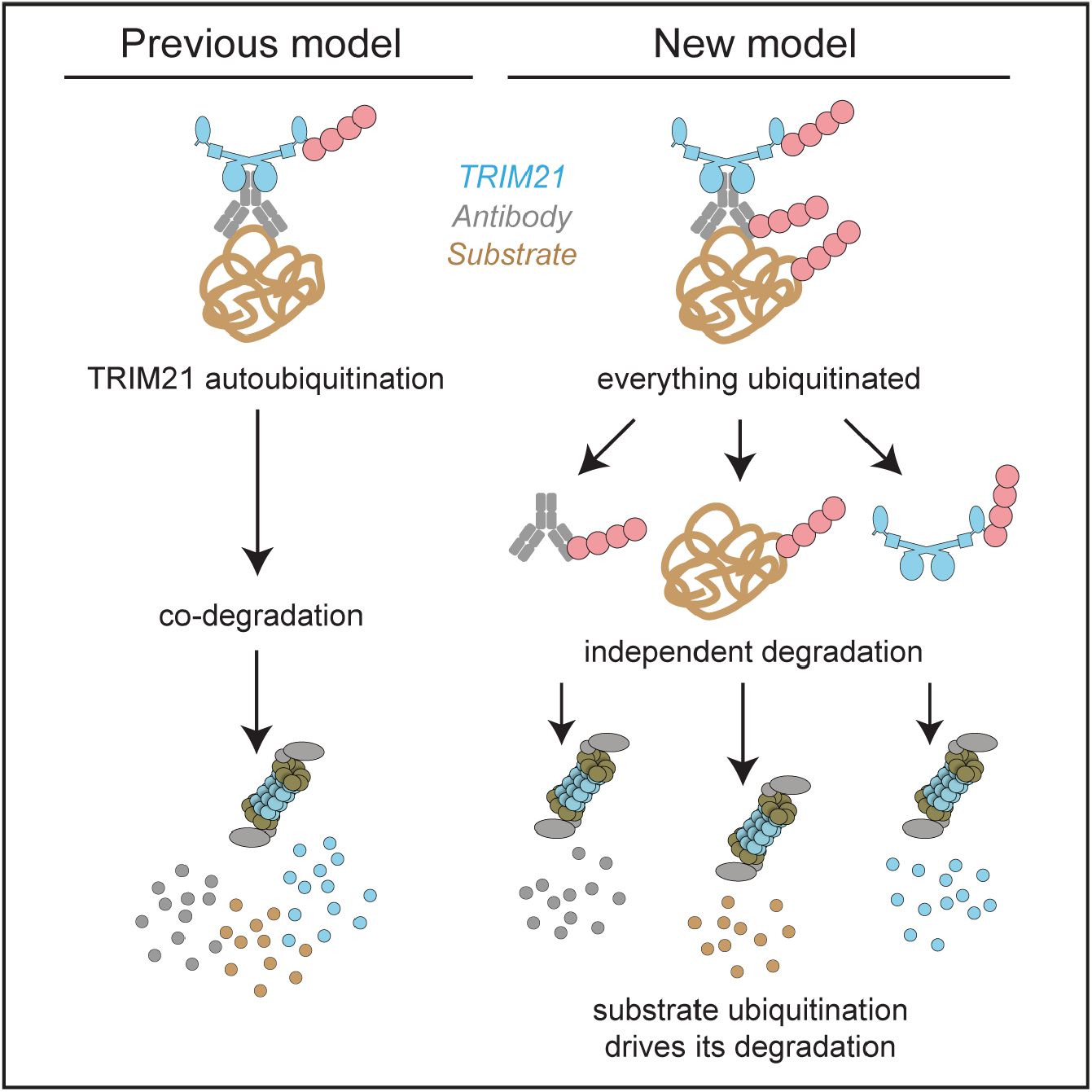
New model for substrate degradation by Trim-Away. See discussion for details.

**Supplementary Table 1:** Data collection and refinement statistics. The TRIM21 RING:Ube2W complex was deposited in the PDB with code 6R6Q. Statistics for both data integration and model refinement are given.

|  | T21-R:Ube2W |
| --- | --- |
| **Data collection** |  |
| Space group | P121 |
| Cell dimensions |  |
| $a, b, c$ (Å) | 62.82, 75.482, 63.835 |
| $\alpha, \beta, \gamma$ (°) | 90.0, 119.307, 90.0 |
| Resolution (Å) | 29.46-2.25 (2.33-2.25) |
| $R_{\text{meas}}$ | 4.8 (65.9) |
| $CC_{1/2}$ (%) | 99.9 (85.9) |
| $I / \sigma I$ | 16.49 (2.01) |
| Completeness (%) | 95.5 (96.3) |
| Redundancy | 3.3 (3.6) |
| **Refinement** |  |
| Resolution (Å) | 29.46-2.25 (2.33-2.25) |
| No. reflections | 23634 (2355) |
| $R_{\text{work}} / R_{\text{free}}$ | 0.22/0.26 (0.36/0.42) |
| No. atoms | 3515 |
| Protein | 3476 |
| Ligand/ion | 4 |
| Water | 35 |
| $B$-factors | |
| Protein | 83.06 |
| Ligand/ion | 56.95 |
| Water | 78.61 |
| R.m.s. deviations |  |
| Bond lengths (Å) | 0.011 |
| Bond angles (°) | 1.41 |

\*Values in parentheses are for highest-resolution shell.

**Supplementary Table 2: Reagents used in this study**. List and details of **(A)** plasmids, **(B)** purified proteins, **(C)** cell lines, **(D)** antibodies and **(E)** commercial assays and software used in this study. See separate excel file.

## METHODS

### Plasmids

A full list of plasmids used in this study, including primary sequences of all constructs can be found in Supplementary Table 2A. For lentivirus production, constructs were inserted into a modified version of pSMPP (Addgene #104970) where the SFFV promotor and puromycin resistance sequences were replaced with PGK1 promoter and Zeocin resistance sequences respectively (pPMEZ). For *in vitro* mRNA transcription, constructs were inserted into pGEMHE^59^, which contains UTR and polyA sequences for optimal mRNA stability and translation. For protein purification, constructs were inserted into derivations of the pOP and pET (Novagen) series of vectors.

### Lentivirus production

Lentivirus particles were collected from HEK293T cell supernatant 3 days after co-transfection (FuGENE 6, Promega) of lentiviral plasmid constructs (Supplementary Table 2A) with HIV-1 GagPol expresser pcRV1 (a gift from Dr. Stuart Neil) and pMD2G, a gift from Didier Trono (Addgene plasmid #12259). Supernatant was filtered at 0.45 µm before storage at −80°C.

### *In vitro* transcription of mRNA

pGEMHE plasmid constructs (Supplementary Table 2A) were linearized and 5’-capped mRNA was synthesized with T7 polymerase (NEB HiScribeT7 ARCA kit) according to manufacturer’s instructions. mRNA concentration was quantified using a Qubit 4 fluorometer (ThermoFisher) and RNA Broad Range assay kit (ThermoFisher; Q10211).

### Protein Expression and purification

A full list of purified proteins used in this study can be found in (Supplementary Table 2B). Ube2W, Ube2N and Ube2V2, and TRIM21 R, R-PS, R-R-PS, T21R-vhhGFP4 and mEGFP were expressed in *Escherichia coli* BL21 DE3. Ubiquitin and Ube1 were expressed in *E. coli* Rosetta 2 DE3 cells as previously described^38^. Cells were grown at 37 °C and 220 rpm until an OD^600^ of ∼0.7. After induction, the temperature was reduced to 18 °C overnight. For TRIM21 and E2s induction was performed with 0.5 mM IPTG and 10 µM ZnCl_2_, for ubiquitin and Ube1 with 0.2 mM IPTG. mEGFP was expressed in ZY autoinduction media^60^ at 37 °C and 220 rpm. At OD^600^ of 0.7, the temperature was reduced to 18 °C for expression overnight. After centrifugation, cells were resuspended in 50 mM Tris pH 8.0, 150 mM NaCl, 10 µM ZnCl_2_, 1 mM DTT, 20 % Bugbuster (Novagen) and Complete protease inhibitors (Roche, Switzerland). For His-tagged proteins, 20 mM imidazole was added to the buffer. Lysis was performed by sonication. TRIM21-R-PS and -R-R-PS were expressed with N-terminal GST-SUMO tag and TRIM-R, Ube2W, Ube2V2 and Ube1 were expressed with N-terminal GST-tag followed by a TEV protease cleavage site and purified via glutathione sepharose resin (GE Healthcare) equilibrated in 50 mM Tris pH 8.0, 150 mM NaCl and 1 mM DTT. The tag was cleaved on beads overnight at 4 °C (with SUMO or TEV protease, respectively). Cleavage with SUMO protease resulted in no cleavage scar on TRIM21-R-PS and -R-R-PS. TEV cleavage results in an N-terminal GSH-scar on TRIM21-R, an N-terminal G-scar on Ube2N, an N-terminal GSQEF-scar on Ube2V2 and an N-terminal GSH-scar on Ube2W. In the case of Ube1, no protease cleavage was performed and the GST-Ube1 fusion protein was eluted using 50 mM Tris pH 8.0, 150 mM NaCl, 10 mM reduced glutathione and 1 mM DTT. mEGFP was expressed with an N-terminal His-tag without protease cleavage site and Ube2N was expressed with an N-terminal His-tag followed by a TEV protease cleavage site. TRIM21 R-vhhGFP4 was expressed as a His-SUMO fusion protein, to generate the native TRIM21 N-terminus after SUMO protease cleavage during purification. His-tagged proteins were purified via Ni-NTA resin equilibrated in 50 mM Tris pH 8.0, 150 mM NaCl, 20 mM imidazole and 1 mM DTT. Proteins were eluted in 50 mM Tris pH 8.0, 150 mM NaCl, 1 mM DTT, and 300 mM imidazole. For Ube2N, TEV-cleavage of the His-tag was performed overnight by dialyzing the sample against 50 mM Tris pH 8.0, 150 mM NaCl, 1 mM DTT, and 20 mM imidazole. Afterward, His-tagged TEV protease was removed by Ni-NTA resin. For T21R-vhhGFP4, SUMO protease cleavage was performed on the Ni-NTA resin overnight at 4 °C. Elution was performed on the next day using the equilibration buffer. Finally, size-exclusion chromatography of all proteins was carried out on either HiLoad 26/60 or 16/600 Superdex 75 prep grade column (GE Healthcare) in 20 mM Tris pH 8.0, 150 mM NaCl, and 1 mM DTT, except for GST-Ube1 for which either HiLoad 26/60 or 16/600 Superdex 200 prep grade column (GE Healthcare) were used. Ubiquitin purification was performed following the protocol established by the Pickart lab^61^. After cell lysis by sonication (lysis buffer: 50 mM Tris pH 7.4, 1 mg mL^-1^ Lysozyme (by Sigma Aldrich, St. Louis, USA), 0.1 mg mL^-1^ DNAse (by Sigma Aldrich, St. Louis, USA), a total concentration of 0.5 % perchloric acid was added to the stirring lysate at 4 °C. The (milky) lysate was incubated for another 30 min on a stirrer at 4 °C to complete precipitation. Next, the lysate was centrifuged (19,500 rpm) for 30 min at 4 °C. The supernatant was dyalized overnight (3,500 MWCO) against 3 L 50 mM sodium acetate pH 4.5. Afterwards, Ub was purified via cation-exchange chromatography using a 20 mL SP column (GE Healthcare) using a NaCl gradient (0 – 1000 mM NaCl in 50 mM NaAc pH 4.5). Finally, size exclusion chromatography was carried out on a HiLoad 26/60 Superdex 75 prep grade column (GE Healthcare) in 20 mM Tris pH 7.4. Isotopically labelled proteins were expressed using *Escherichia coli* BL21 DE3 cells in M9 minimal media supplemented with ^15^NH_4_Cl (Sigma-Aldrich ISOTEC). *Chaetomium thermophilum* Naa50^82-289^ containing a C-terminal His-tag was expressed using *E. coli* Rosetta 2 cells in ZY autoinduction media^60^ which was grown at 37 °C and 220 rpm. At OD^600^ of 0.7, the temperature was reduced to 18 °C for expression overnight. *Ct*Naa50^82-289^ was purified as follows: Cells were harvested, resuspended in buffer A500 (20 mM HEPES pH 7.5, 500 mM NaCl, 20 mM imidazole) supplemented with a protease inhibitor mix (SERVA Electrophoresis GmbH, Germany) and lysed with a microfluidizer (M1-10L, Microfluidics). The lysate was cleared for 30 min at 50,000 g, 4 °C and filtered through a 0.45 µm membrane. The supernatant was applied to a 1 mL HisTrap FF column (GE Healthcare) for Ni-IMAC (immobilized metal affinity chromatography) purification. The column was washed with buffer A500 and the proteins were eluted with buffer A500 supplemented with 250 mM imidazole. *Ct*Naa50^82-289^ was subsequently purified by SEC (size-exclusion chromatography) using a Superdex 75 26/60 gel filtration column (GE Healthcare) in buffer G500 (20 mM HEPES pH 7.5, 500 mM NaCl). SUMO protease (MBP-Ulp1 (based on R3 sequence^62^) was purified using an MBPTrap HP 5 ml column and eluted with 50 mM Tris pH 8, 150 mM NaCl, 1 mM DTT and 10 mM Maltose. Finally, the eluted fractions were separated on a HiLoad 26/60 Superdex 75pg SEC column (150 mM NaCl, 50 mM Tris pH 8 and 1 mM DTT).

### Cell culture

HEK293T (ATCC) and NIH3T3-CAV1-EGFP^63^ cells were cultured in DMEM medium (Gibco; 31966021) supplemented with 10% calf serum and penicillin-streptomycin. RPE-1 cells (ATCC) were cultured in DMEM/F-12 medium (Gibco; 10565018) supplemented with 10% Calf Serum and penicillin-streptomycin. All cells were grown at 37°C in a 5% CO_2_ humidified atmosphere and regularly checked to be mycoplasma-free. The sex of NIH3T3 cells is male. The sex of HEK293T and RPE-1 cells is female. For proteasome inhibition experiments, MG132 (Sigma; C2211) was used at a final concentration of 25 µM and Epoxomicin (Sigma; 324801) was used at 10 µM. Following electroporation, cells were grown in medium supplemented with 10% calf serum without antibiotics. Live imaging was performed using the IncuCyte S3 live cell analysis system (Sartorius) housed within a 37°C, 5% CO_2_ humidified incubator. For live imaging with the IncuCyte, cell culture medium was replaced with Fluorobrite (Gibco; A1896701) supplemented with 10% calf serum and GlutaMAX (Gibco; 35050061).

### Cell lines

Cell lines used and generated in this manuscript are detailed in (Supplementary Table 2C). RPE-1 TRIM21 KO cells^15^, HEK293T TRIM21 KO cells^64^ and NIH3T3-CAV1-EGFP^63^ were described previously. For stable expression of CAV1-mEGFP and CAV1-mEGFP-Halo, RPE-1 cells were transduced with lentiviral particles at multiplicity ∼0.1 transducing units per cell and the GFP-positive population selected by flow cytometry. For stable expression of TRIM21-HA at endogenous levels, RPE-1 TRIM21 KO cells were reconstituted with TRIM21-HA construct under control of the native TRIM21 promoter as described previously^64^.

### Electroporation

Electroporation was performed using the Neon® Transfection System (Thermo Fisher). Cells were washed with PBS and resuspended in Buffer R at a concentration of between 1-8 × 10^7^ cells ml^-1^. For each electroporation reaction 1 - 8 × 10^5^ cells in a 10.5 µl volume were mixed with 2 µl of antibody (typically 0.5 mg/ml) or mRNA (typically 0.5 µM) or protein to be delivered. The mixture was taken up into a 10 µl Neon® Pipette Tip, electroporated at 1400V, 20 ms, 2 pulses and transferred to media without antibiotics.

### Measurement of fluorescence in live cells

To quantify GFP fluorescence in live cells, images were acquired and analysed using the IncuCyte live cell analysis system (Sartorius). Within the IncuCyte software, the integrated density (the product of the area and mean intensity) for GFP fluorescence was normalized to total cell area (phase) for each image. Values were normalized to internal controls within each experiment.

### Antibodies

Antibodies and concentrations used for traditional immunoblotting (IB), capillary-based immunoblotting (Jess) and electroporation (EP) are detailed in (Supplementary Table 2D). All antibodies used for electroporation were either purchased in azide-free formats or passed through Amicon Ultra-0.5 100 KDa centrifugal filter devices (Millipore) to remove traces of azide and replace buffer with PBS.

### Adv5 neutralization assay

Adenovirus serotype 5 2.6-del CMV-eGFP (Adv5-GFP, Viraquest) was diluted to 1.1×10^9^ T.U./mL in PBS, and 16 uL was incubated 1:1 with the anti-hexon reconmbinant humanized IgG1 9C12 or 9C12^H433A 65^ at indicated concentrations, or PBS. After 1 hour incubation at room temperature, complexes were diluted with 250 µL Fluorobrite media and used for Adv5 neutralization assays. For infections, 4×10^6^ HEK293T TRIM21 KO cells were electroporated with PBS or R-R-PS ± N^α^-acetylation and resuspended in 2 mL Fluorobrite media. 50 µL of each cell suspension was combined 1:1 with Adv5:9C12 or Adv5:9C12^H433A^ complexes (for immediate infections) or Fluorobrite (for delayed infections) in 96-well plates. For delayed infections, electroporated cells were allowed to adhere to the plate for 2 hours, then media was replaced with 50 µL of Fluorobrite and infected with 50 µL Ad5:9C12 complexes. Infection levels were quantified using the IncuCyte system by measuring GFP fluorescence area relative to total cell area 16h post-infection. Infection levels are plotted relative to Adv5-GFP infection the absence of 9C12 antibody.

### NFκB signalling assay

HEK293T TRIM21 KO cells were transfected with 2ug pGL4.32 NF-κB luciferase plasmid (Promega), using 12 uL of Viafect (Promega) in 200 uL OptiMEM (Thermo Fisher). Twenty-four hours later 4×10^6^ transfected cells were electroporated with PBS or R-R-PS ± N^α^-acetylation and resuspended in 1 mL DMEM media. For infections, Adv5:9C12 complexes were prepared as described above, except Ad5-GFP was diluted to 1.1×10^10^, 9C12 was used at 20 ug/mL, and the complex was diluted into 150 µL DMEM after 1 hour incubation. 50 µL of the electroporated cell suspension was mixed 1:1 with Adv5:9C12 complexes or PBS (control), then lysed 4 hours later in 100 uL of SteadyLite Plus luciferase reporter (PerkinElmer). As an internal control, TNF-α was used at 10 ng/uL. Luminance was recorded on a PheraStar FS (BMG LabTech).

### Immunoblotting

Samples were run on NuPAGE 4-12% Bis-Tris gels (ThermoFisher) and transferred onto nitrocellulose membrane. Membranes were incubated in blocking buffer (PBS, 0.1% Tween20, 5% milk) for 1h at room temp prior to incubation with antibodies. Antibodies and dilutions (in blocking buffer) used for immunoblotting (IB) are detailed in (Supplementary Table 2D). HRP-coupled antibodies were detected by enhanced chemiluminescence (Amersham, GE Healthcare) and X-ray films. IRDye-coupled antibodies were detected using LI-COR Odyssey CLx imaging system.

### Capillary-based immunoblotting

RIPA buffer protein extracts were diluted 1:2 in 0.1x sample buffer (bio-techne; 042-195) and run on the Jess Simple Western system using a 12-230kDa separation module (bio-techne) according to manufacturer’s instructions. Antibodies and dilutions used for capillary-based immunoblotting (Jess) are detailed in (Supplementary Table 2D). Protein peak areas were quantified using Compass software (bio-techne) and normalized to internal protein loading controls within each capillary.

### In vitro ubiquitination assays

Ube2W-dependent TRIM21-mono-ubiquitination assays were performed in 50 mM Tris pH 7.4, 150 mM NaCl, 2.5 mM MgCl_2_ and 0.5 mM DTT. The reaction components were 2 mM ATP, 1 µM GST-Ube1, 80 µM ubiquitin and the indicated concentrations of Ube2W and TRIM21, respectively. The reaction was stopped by addition of LDS sample buffer containing 50 mM DTT at 4 °C. Next, samples were boiled at 90 °C for 2 min. For reactions using 10 µM TRIM21, visualization was performed by Instant Blue stained LDS-PAGE only. Polyubiquitin chain extension assays were performed as above, but instead of Ube2W, 0.5 µM Ube2N/Ube2V2 were added. For antibody-induced mono-ubiquitination similar conditions were used as for the LDS-PAGE analysed mono-ubiquitination described above. However, the concentration of TRIM21 was reduced to 100 nM and GST-Ube1 to 0.25 µM. Anti-GFP antibody (9F9.F9) was added in one molar equivalent to TRIM21. The reaction was initiated by addition of Ube2W (0, 50, 100, 200 nM). The reaction was stopped by addition of LDS sample buffer at 4 °C. Samples were boiled at 90 °C for 2 min and resolved by LDS-PAGE. TRIM21 was visualized using western blot. In vitro reconstitution of Trim-Away ubiquitination events was performed similar to the antibody-induced mono-ubiquitination experiments described above. E2 concentrations were 200 nM Ube2W and 0.5 µM Ube2N/Ube2V2 and His-mEGFP was used as Trim-Away target at 200 nM.

### Acetylation and monoubiquitination assay

N-terminal acetylation of TRIM21 was introduced by the *Chaetomium thermophilum* N-acetyl transferase (NAT) Naa50 catalytic domain residues 82 to 289 (*Ct*Naa50^82-289^). Acetylation reactions were performed in 50 mM Tris pH 7.4 and 150 mM NaCl for 4 h at 25 °C. The reactions contained 20 µM TRIM21, 1 mM Acetyl-CoA and 1 µM *Ct*Naa50. After the Acetylation reaction was finished, it was mixed 1:1 with a Ube2W-ubiquitination mix containing 100 mM Tris pH 7.4, 300 mM NaCl, 5 mM MgCl_2_ and 1 mM DTT, 4 mM ATP, 2 µM GST-Ube1, 160 µM ubiquitin and 2 µM Ube2W. The Ube2W ubiquitinaton reaction was performed for 1 h at 37 °C and stopped by addition of LDS sample buffer containing 50 mM DTT at 4 °C, followed by boiling the samples at 90 °C for 2 min. Visualization was performed by Instant Blue stained LDS-PAGE only.

### NMR spectroscopy

Two-dimensional NMR measurements (^15^N-HSQC) were performed at 25 °C on Bruker Avance I 600 MHz spectrometer equipped with 5mm ^1^H-^13^C-^15^N cryogenic probe. Data was processed with the program Topspin (Bruker BioSpin GmbH, Germany) and analyzed with the program CCPN analysis v2^66^. Samples were buffer exchanged into 50 mM deuterated Tris pH 7.0, 150 mM NaCl and 1 mM deuterated DTT (Cambridge Isotopes, United Kingdom).

Chemical shift perturbations (CSPs) were calculated using the equation (3):

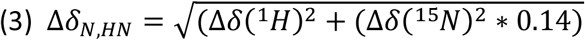

where Δd_N, HN_ is the CSP, Δδ(^1^H) and Δδ(^15^N) are the chemical shift differences between the position of proton or nitrogen signal in absence and presence of titrant. TRIM21 assignments were used from a previous publication^14^.

### Mass spectrometry

Excised protein gel pieces were destained with 50 % v/v acetonitrile: 50 mM ammonium bicarbonate. After reduction with 10 mM DTT and alkylation with 55 mM iodoacetamide, the proteins were digested overnight at 37 °C with 6 ng μL^-1^ of Asp-N (Promega, UK). Peptides were extracted in 2 % v/v formic acid : 2 % v/v acetonitrile and subsequently analyzed by nano-scale capillary LC-MS/MS with an Ultimate U3000 HPLC (Thermo Scientific Dionex, San Jose, USA) set to a flowrate of 300 nL min^-1^. Peptides were trapped on a C18 Acclaim PepMap100 5 μm, 100 μm × 20 mm nanoViper (Thermo Scientific Dionex, San Jose, USA) prior to separation on a C18 T3 1.8 μm, 75 μm × 250 mm nanoEase column (Waters, Manchester, UK). A gradient of acetonitrile eluted the peptides, and the analytical column outlet was directly interfaced using a nano-flow electrospray ionization source, with a quadrupole Orbitrap mass spectrometer (Q-Exactive HFX, ThermoScientific, USA). For data-dependent analysis a resolution of 60,000 for the full MS spectrum was used, followed by twelve MS/MS. MS spectra were collected over a *m*/*z* range of 300–1,800. The resultant LC-MS/MS spectra were searched against a protein database (UniProt KB) using the Mascot search engine program. Database search parameters were restricted to a precursor ion tolerance of 5 ppm with a fragmented ion tolerance of 0.1 Da. Multiple modifications were set in the search parameters: two missed enzyme cleavages, variable modifications for methionine oxidation, cysteine carbamidomethylation, pyroglutamic acid and protein N-term acetylation. The proteomics software Scaffold 4 was used to visualize the fragmented spectra.

### Statistical Analysis

Average (mean), standard deviation (s.d.), standard error of the mean (s.e.m) and statistical significance based on Student’s *t*-test (two-tailed) and one- or two-way ANOVA were calculated in Microsoft Excel or Graphpad Prism. Significance are represented with labels ns (not significant, P>0.05), * (P≤0.05), ** (P≤0.01), *** (P≤0.001), **** (P≤0.0001).

### Crystallography

Crystals of TRIM21-RING:Ube2W^V30K/D67K/C91K^ complex were grown in 2 nl drops at 10 mg/ml at 17 °C in 0.1 M Bicine pH 9.0, 5% PEG 6000, 0.1M TCEP hydrochloride. Diffraction experiments were performed at the European Synchrotron Radiation Facility at beamline ID23 using a Dectris PILATUS 6M detector at a wavelength of 0.984004 Å. The diffraction data at 2.25 Å was processed using XDS. The structure was solved by molecular replacement using Phaser^67^ with TRIM21 RING domain (5OLM^14^) and Ube2W residues 1-118 (2MT6^68^) as search models. Model building and real-space-refinement were carried out in coot^69^, and refinement was performed using REFMAC5 and phenix-refine^70^. For full data collection and refinement statistics see Supplementary Table 1. Model and structure factors have been deposited at the PDB with the accession code 8A58.

